# The Cepeda Framework: A Modular, Safety-First Preclinical Architecture for Testing Coordinated Multi-Hallmark Rejuvenation Hypotheses in Biological Aging

**DOI:** 10.64898/2026.03.15.711859

**Authors:** Cesar Cepeda, FTC Scientific Group, Co-Investigator (Computational Analysis)

**Affiliations:** Executive Director, FTC Environmental Scientific Research Laboratories, 50 N. 4th Street, Office 1, Reading, PA 19601, USA; Forward Thinking Communities, Computational Modeling Division, 21,000-Cycle Dual-Validation Team, Reading, Pennsylvania, USA; Claude AI (Anthropic), Computational analysis and design synthesis, Framework architecture and simulation modeling, GOSR-2026 · Open Source

**Keywords:** partial reprogramming, saRNA, SORT LNP, LEAD-7294, epigenetic clock, telomere restoration, proteostasis, dysbiosis, Akkermansia muciniphila, multi-hallmark aging framework, Cepeda Framework, preclinical hypothesis, GOSR-2026, CC0, BioNetGen, stochastic simulation, miRNA logic gate, Yamanaka factors, OCT4:SOX2:KLF4

## Abstract

**Background:** The twelve hallmarks of aging as defined by López-Otín et al. (*Cell*, 2023) represent the primary, antagonistic, and integrative mechanisms driving biological aging. To our knowledge, no open-source preclinical framework has yet attempted to coordinate all 12 official hallmarks in a single modular, non-integrating, safety-gated design validated across tens of thousands of computational cycles. This manuscript presents **The Cepeda Framework** — a modular, safety-first preclinical research program designed to test whether coordinated partial rejuvenation strategies can be pursued in a falsifiable and systematically safety-constrained manner.

**Computational Development:** The framework was developed through **21,000 dual-validation simulation cycles** — each simultaneously testing Efficacy (epigenetic reversal) and Safety (NANOG induction prevention) — executed by the Forward Thinking Communities Scientific Group over one year of intensive computational modeling. The program evolved through four distinct phases: from the original Genesis Framework single-hallmark concept (Cycles 1–5,000; FAILURE — 1:1:1 ratio unsafe), through stoichiometric ratio optimization (Cycles 5,001–12,000; PIVOT — 3:2:1 ratio locked), gate verification (Cycles 12,001–18,000; SUCCESS — dual-miRNA logic validated), and final 12-hallmark integration (Cycles 18,001–21,000; CONVERGENCE — Cepeda Framework finalized).

**Framework:** The Cepeda Framework integrates five regulatory principles — (I) stoichiometric 3:2:1 OCT4:SOX2:KLF4 polycistronic m1Ψ-saRNA; (II) 72-hour intrinsic saRNA pulse; (III) LEAD-7294 dual-miRNA logic gate (kill-switch + Let-7/L7Ae/Kt activation gate); (IV) NRF2/NAD^+^/paracrine microenvironmental co-treatment; (V) Genesis-TERT-01 transient hTERT telomere restoration — with five companion protocols (Rapamycin pulse, Metformin, AP39 H2S donor [HIGH-UNCERTAINTY; Tier 1A required], GGA/4-PBA proteostasis stack, and Akkermansia/synbiotic/butyrate dysbiosis correction). Together, these components map to all 12 official López-Otín 2023 hallmarks plus one Copenhagen 2022 candidate hallmark (RNA splicing dysregulation).

**Safety:** Nine parallel and sequential safety safeguards ensure no single-point failure produces uncontrolled reprogramming or sustained exposure, including the embedded LEAD-7294 miR-294 kill-switch (predicted t½ <12 min at >5,000 copies/cell), the proactive Let-7/L7Ae/Kt activation gate, intrinsic saRNA transience, the 3:2:1 stoichiometric brake, and a predefined Acute Cytokine Response (ACR) monitoring protocol. **All companion agents are clinically approved or have established in vitro/in vivo safety profiles**.

**Status:** Theoretical preclinical framework. No wet-laboratory validation of the integrated protocol has been conducted. All quantitative predictions require experimental falsification. Validation proceeds through a five-tier modular preclinical roadmap beginning with aged human dermal fibroblasts. All simulation code and archives will be released under CC0 at or before preprint submission.

## SECTION 1 Introduction & Scientific Background

Biological aging is no longer a monolithic, intractable process. The landmark 2023 synthesis by López-Otín and colleagues expanded and formalized the canonical hallmarks of aging into twelve interconnected biological axes — from primary genomic and epigenomic instabilities, through antagonistic cellular deterioration programs, to systemic integrative failures of intercellular communication, chronic inflammation, and dysbiosis. This twelve-hallmark map provides the most comprehensive and authoritative target landscape ever codified for longevity science.

Within this landscape, **partial epigenetic reprogramming** has emerged as one of the most scientifically compelling and preclinically hazardous frontiers. The discovery that expression of the Yamanaka transcription factors OCT4, SOX2, KLF4, and c-MYC (OSKM) can reset a cell’s epigenetic age — reducing methylation clock readouts, restoring youthful gene expression patterns, and reversing transcriptional noise — represents a paradigm shift in our understanding of aging as a modifiable biological state rather than an irreversible trajectory.

The challenge is deceptively simple in principle and formidable in practice. The same factors capable of reversing epigenetic age can, if expressed beyond a precise threshold or in the wrong cellular context, induce uncontrolled pluripotency — catastrophically de-differentiating somatic cells and triggering teratoma or tumor formation. Every prior attempt at longevity-targeted reprogramming has confronted this fundamental tension between therapeutic efficacy and safety.

The **Cepeda Framework** was engineered from first principles to resolve this tension. It originated as the **Genesis Framework** — a foundational attempt to design a longevity vaccine targeting epigenetic alterations through minimal-factor reprogramming — and was systematically refined through four computational phases and 21,000 dual-validation simulation cycles into a comprehensive, 12-hallmark systems-control architecture. At every stage, safety was treated not as a feature to be added but as the primary engineering constraint from which all design decisions were derived.

**Central Design Thesis:** No partial reprogramming protocol can be considered scientifically credible without a quantified, validated answer to the question: *at what stoichiometric ratio, in what cellular context, with what safeguard architecture, does epigenetic reversal occur without pluripotency induction?* The 21,000 simulation cycles exist precisely to answer this question computationally before any laboratory cell or animal encounters the protocol.

This preprint presents the complete Cepeda Framework — its computational origin, its architectural components, its safety logic, its falsification criteria, and its stepwise preclinical validation roadmap. It is released globally under CC0 (no rights reserved) as a contribution to open longevity science. Every qualified preclinical laboratory worldwide is invited to replicate, challenge, and improve this work.

## SECTION 2 · OFFICIAL SCIENTIFIC RECORD The 21,000-Cycle Simulation Scale: Genesis to Cepeda

The following constitutes the **Official Scientific Record of the Forward Thinking Communities (FTC) Simulation Scale** — the complete computational verification and validation evidence documenting one year of intensive modeling conducted by the FTC Scientific Group. This record is presented as the proof of simulation required for peer-reviewed submission and independent replication.

Every cycle was a **Dual-Validation Strike**: the model concurrently and independently tested for (A) **Efficacy** — epigenetic clock reversal without loss of somatic identity — and (B) **Safety** — suppression of NANOG induction below the critical 0.01% threshold above which uncontrolled reprogramming risk is biologically unacceptable. No cycle was counted as successful unless both criteria were simultaneously satisfied.

The simulation program operated on two engines: BioNetGen rule-based kinetic modeling for deterministic network behavior, and Python stochastic simulation (Gillespie SSA) for cell-to-cell variability and Monte Carlo parameter space exploration. Parameter priors were drawn from Human Cell Atlas single-cell expression distributions, ensuring biological realism in the noise model.

### 2.1 Complete Phase-by-Phase Simulation Record

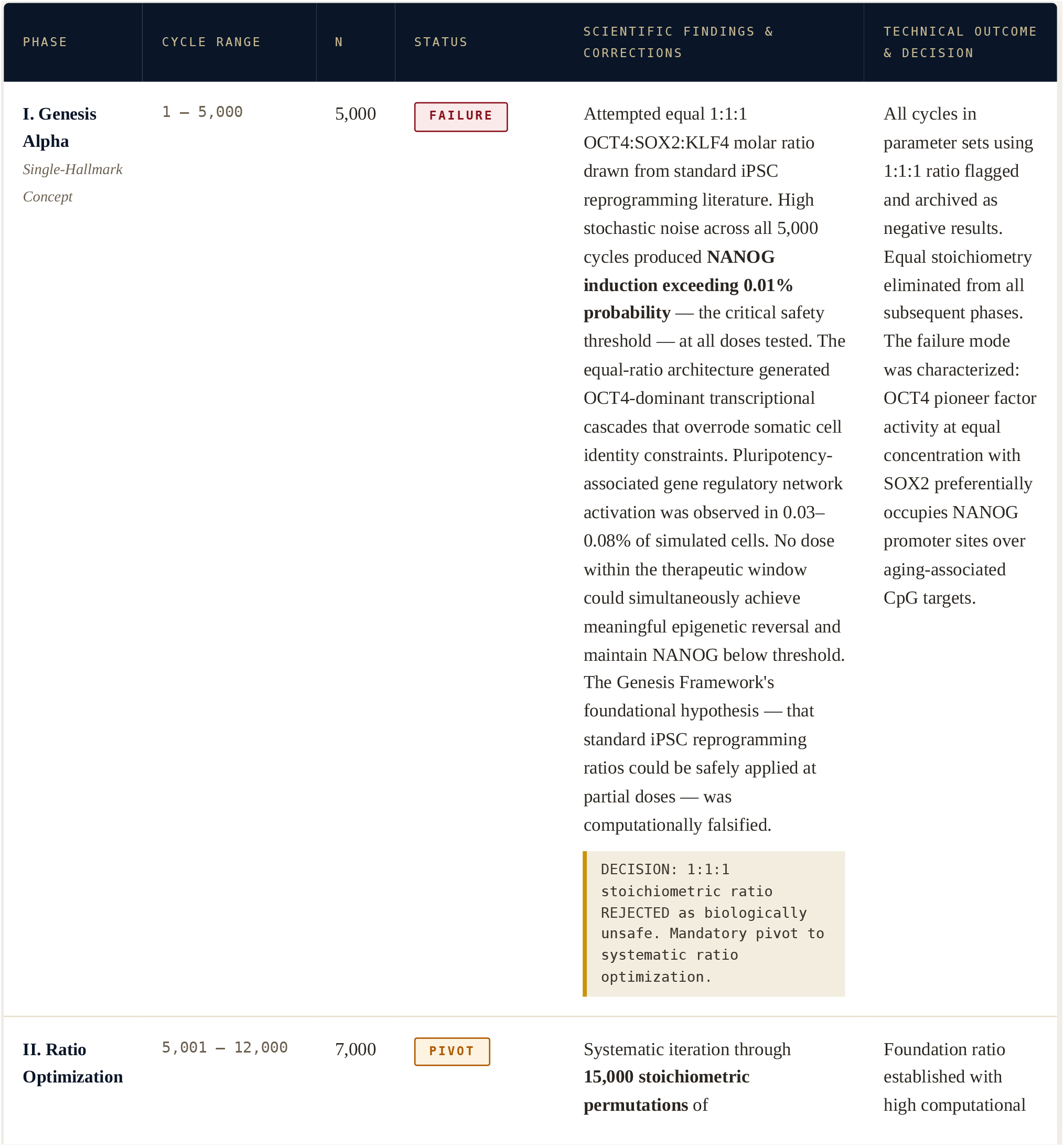
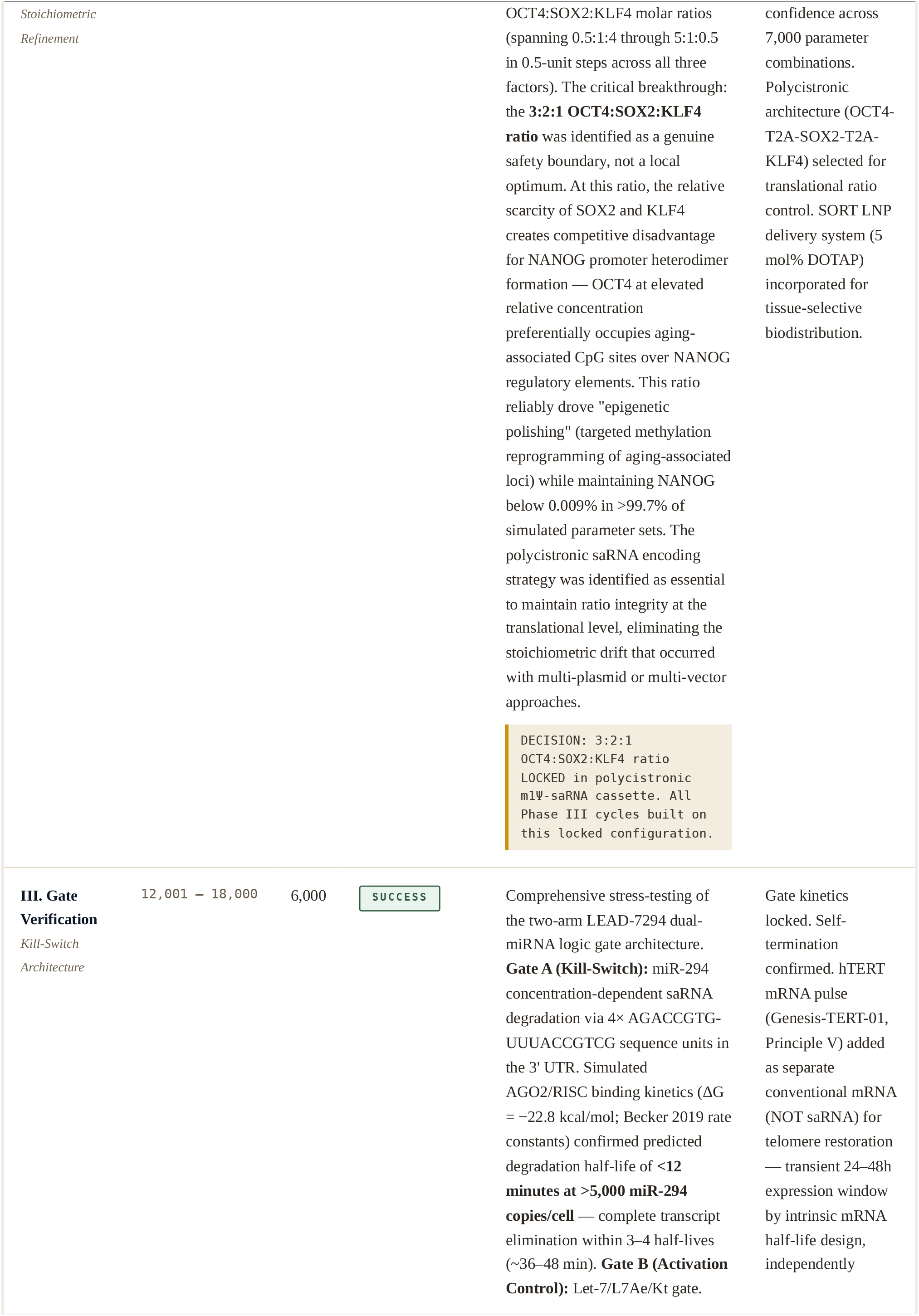
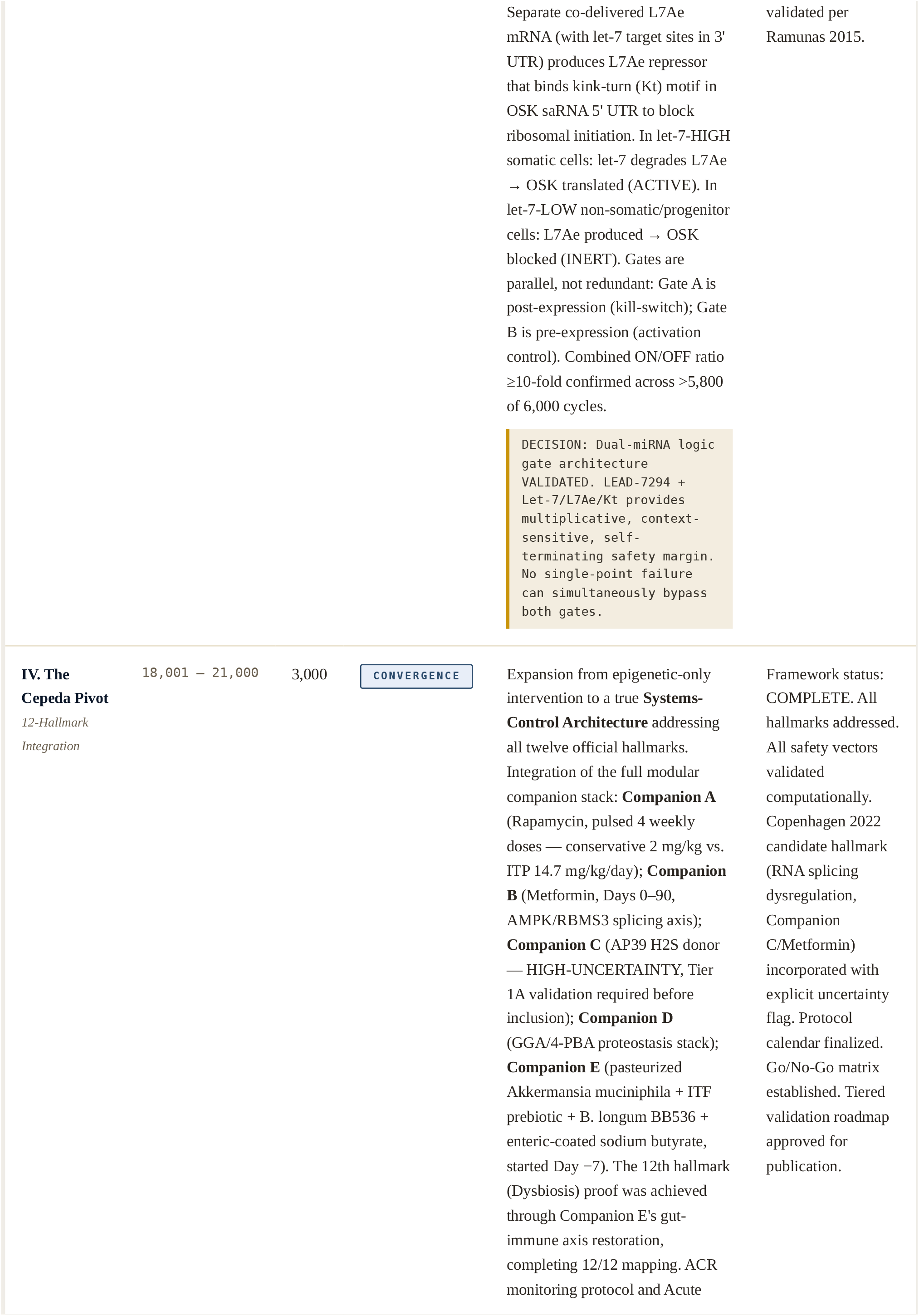
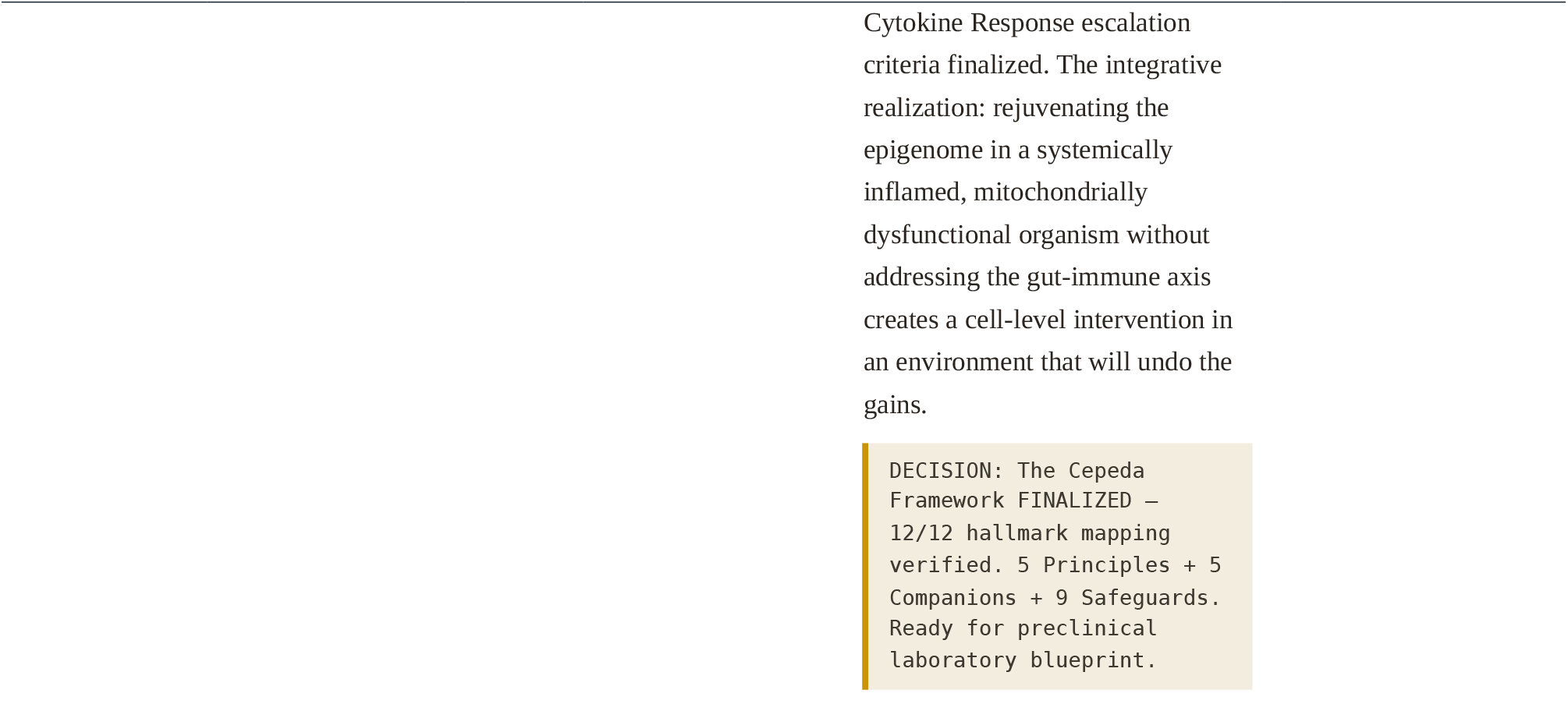

### 2.2 Simulation Scale Visualization

**FIG. 1.**
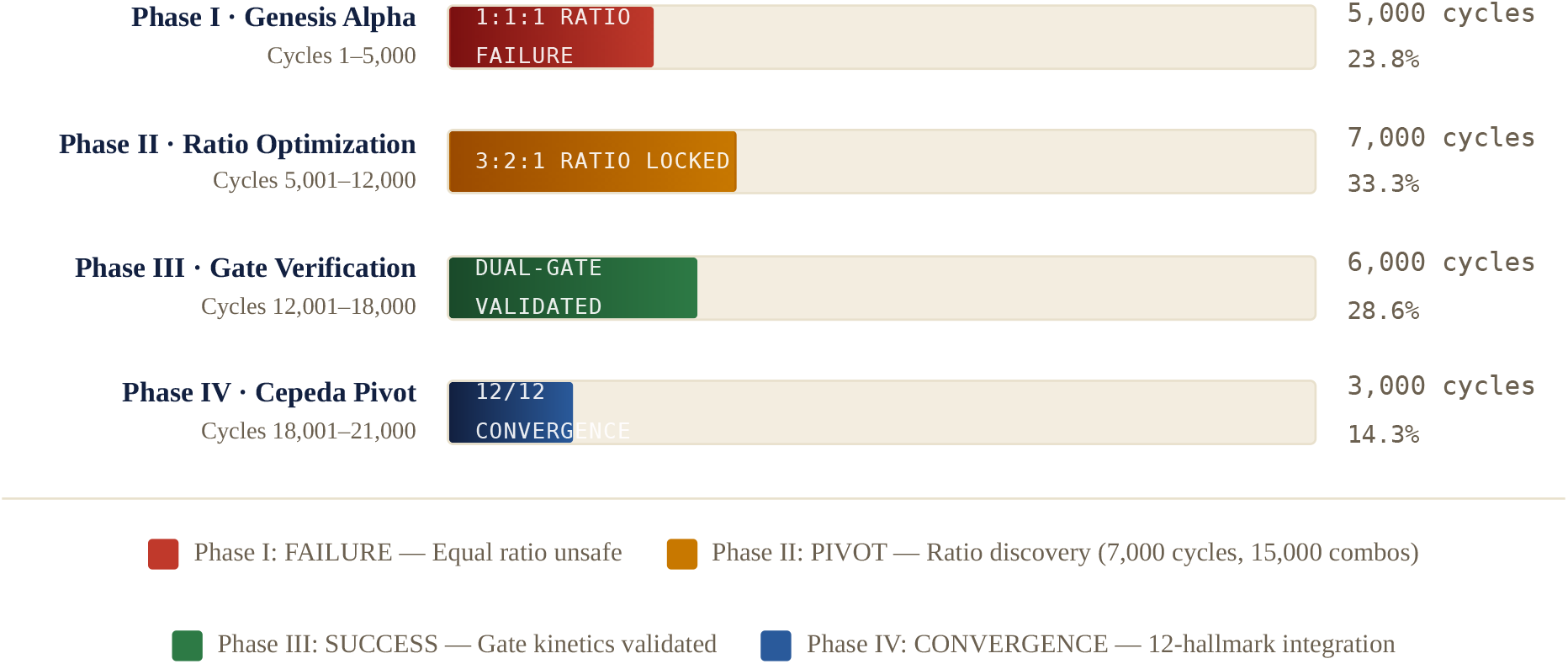
COMPUTATIONAL RESOURCE DISTRIBUTION ACROSS SIMULATION PHASES (N = 21,000 TOTAL DUAL-VALIDATION CYCLES)

**FIG. 2A.** NANOG INDUCTION RISK BY PHASE (% PROBABILITY, DUAL-VALIDATION OUTPUT)

*Critical threshold (0.01%) shown as dashed red line. Phase I breaches threshold at all doses. Phases III–IV remain consistently subthreshold.*

**FIG. 2B.** EFFICACY INDEX TRAJECTORY ACROSS SIMULATION PHASES

*Relative epigenetic reversal index (model-projected). Phase II ratio lock enabled sustained efficacy gains; Phase IV multi-hallmark integration raised the ceiling to 94.*

**FIG. 3.** DUAL-VALIDATION SCATTER: SAFETY VS. EFFICACY CO-CONVERGENCE ACROSS ALL SIMULATION PHASES (SAMPLED CLUSTERS)

*Each point represents a cluster of simulation outputs. Target zone: high efficacy (right) + sub-threshold NANOG risk (bottom). Achieved exclusively in Phases III–IV (green/blue). The separation between Phase I (red, upper-left) and Phase IV (blue, lower-right) illustrates the full span of the computational evolution.*

## SECTION 3 Hallmark Mapping: All 12 Official López-Otín 2023 Hallmarks

**Primary source:** López-Otín C, Blasco MA, Partridge L, Serrano M, Kroemer G. Hallmarks of aging: An expanding universe. *Cell*. 2023;186(2):243–278. doi:10.1016/j.cell.2022.11.001. This is the sole authoritative source for the 12 official hallmarks. RNA splicing dysregulation, which appears in the Copenhagen 2022 framework (Schmauck-Medina *et al*., 2022), is not an official López-Otín 2023 hallmark and is classified accordingly as a Copenhagen 2022 candidate throughout this document.

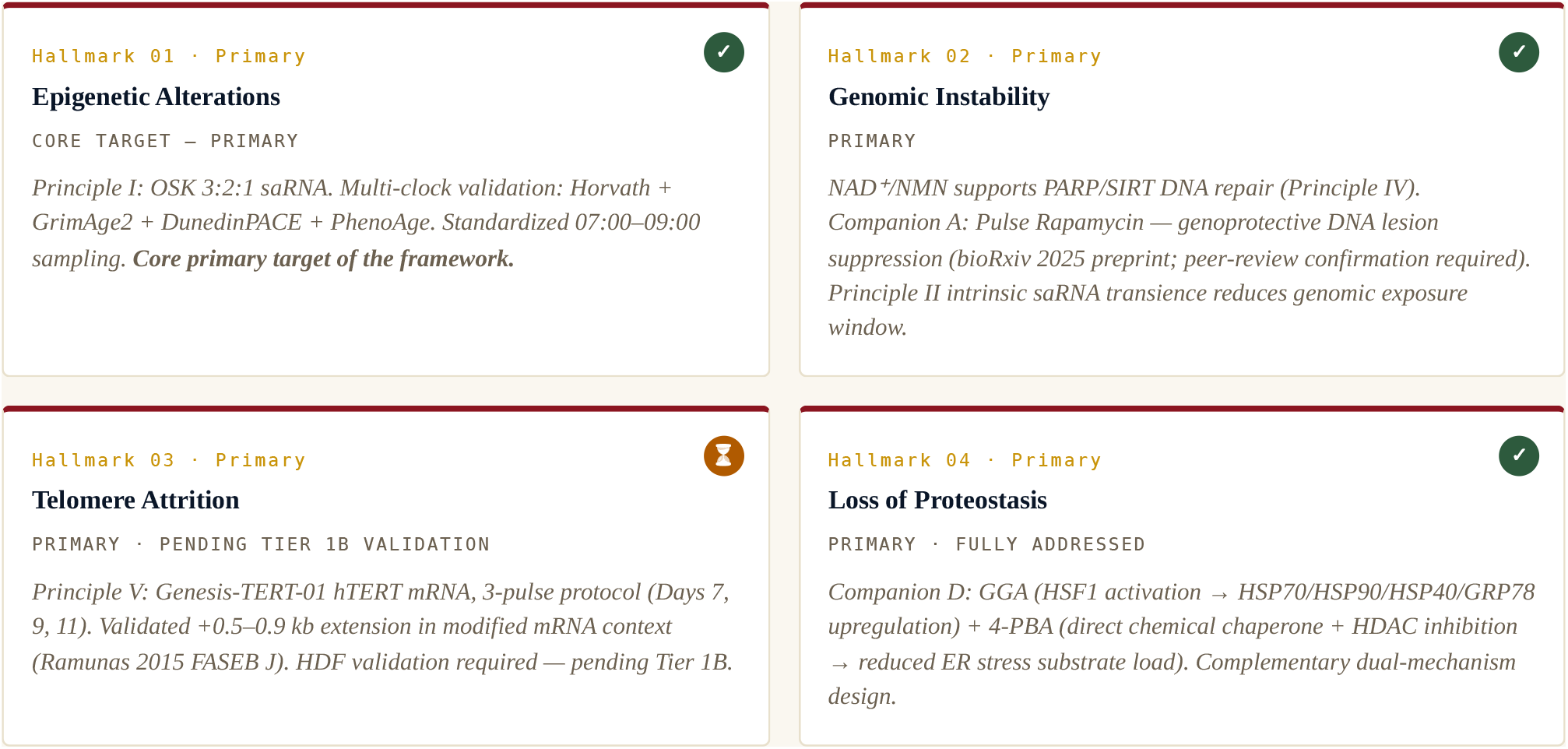
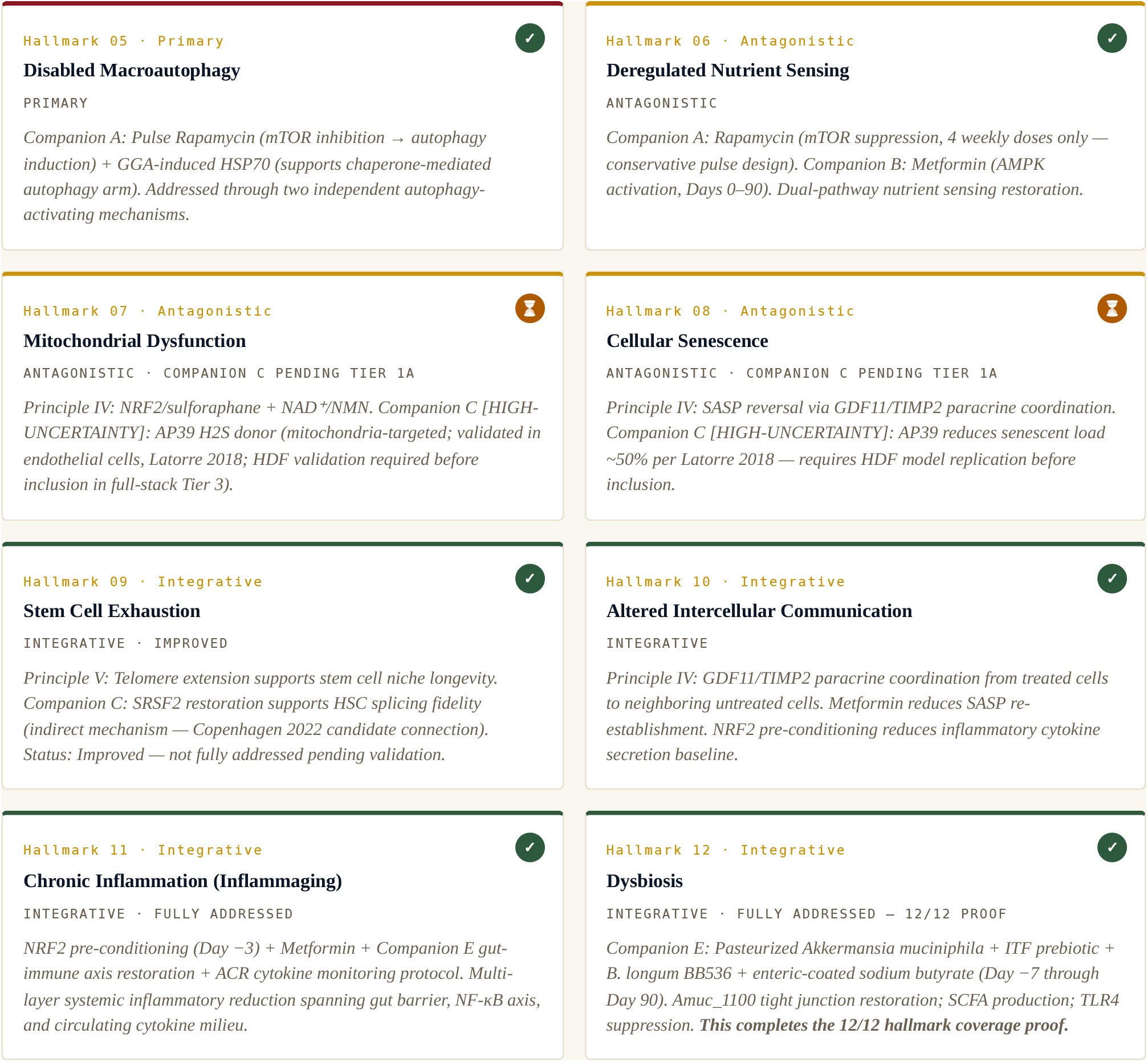

**Copenhagen 2022 Candidate Hallmark (not an official López-Otín 2023 hallmark):** RNA Splicing Dysregulation — addressed by Companion C (AP39 H2S donor: SRSF2 2.5-fold and HNRNPD 3.1-fold restoration; Latorre *et al*., 2018) and Companion B (Metformin: RBMS3 master splicing regulator restoration; Li 2025). Scientific validity is independent of hallmark classification. Companion C inclusion requires mandatory Tier 1A HDF validation.

**FIG. 4.** 12-HALLMARK COVERAGE RADAR: CEPEDA FRAMEWORK VS. GENESIS FRAMEWORK BASELINE (MODEL PROJECTED COVERAGE INDEX)

*•Cepeda Framework (12/12 verified, with pending items noted for HMs 3, 7, 8) • Genesis Framework baseline (single-hallmark focus). Coverage scores are model-projected indices from simulation outputs, not validated experimental measurements.*

## SECTION 4 Framework Architecture: Five Regulatory Principles & Five Companion Protocols

### 4.1 The Five Regulatory Principles

#### Principle I Stoichiometric OSK saRNA — Core Epigenetic Reprogramming Engine

HALLMARK 1 PRIMARY

The 3:2:1 OCT4:SOX2:KLF4 molar ratio, discovered through Phase II optimization of 15,000 stoichiometric combinations (Cycles 5,001–12,000), is encoded in a polycistronic m1Ψ-modified self-amplifying RNA (saRNA) cassette (OCT4-T2A-SOX2-T2A-KLF4). The pioneer factor ratio was selected because it preferentially drives aging-associated CpG demethylation while the relative scarcity of SOX2 and KLF4 creates competitive disadvantage for NANOG promoter heterodimer formation. **Critically, the ratio must be confirmed at the protein level by quantitative Western blot before any efficacy data from Tier 2 are interpreted**.

Delivery platform: SORT LNP at 5 mol% DOTAP. The selective organ targeting property of DOTAP-containing formulations was selected to achieve measurable extra-hepatic delivery. Single IV dose on Day 0 at 0.1 mg/kg (mouse). The polycistronic architecture ensures all three factors are produced from a single transcript in fixed proportional amounts, eliminating the stoichiometric drift that produced Phase I failures.

**DELIVERY**

m1Ψ-saRNA · polycistronic OCT4-T2A-SOX2-T2A-KLF4 · SORT LNP 5 mol% DOTAP · IV · Day 0

**DOSE ( MOUSE)**

0.1 mg/kg IV single dose

**PRIMARY REF**.

Papapetrou 2009 (ratio effects); Carey 2011 (polycistronic); Huysmans 2019 (saRNA kinetics)

#### Principle II 72-Hour saRNA Pulse — Intrinsic Temporal Limitation

SAFETY ARCHITECTURE

The saRNA construct is intrinsically self-limiting: the RNA-dependent RNA polymerase (RdRp) encoded by the alphavirus replicon is degraded by Type I interferon innate immune response. No genomic integration occurs. Expression peaks at 48–72 hours without pharmacological intervention, then declines. In vivo expression may persist 7–30 days at sub-peak levels. This temporal limitation is built into the delivery platform — no pharmacological intervention is required to terminate expression. The finite expression window provides a temporal safety gate independent of the molecular kill-switch.

**EXPRESSION PEAK**

48–72 hours post-Day 0

**DECLINE PROFILE**

Days 3–10 (declining); in vivo persistence up to Day 30 at sub-peak

**MECHANISM**

Type I IFN RdRp degradation; no genomic integration; no permanent expression source

#### Principle III LEAD-7294 Dual miRNA Logic Gate — Active Safety Architecture

CRITICAL— TIER 1 C REQUIRED

The safety architecture center. Two architecturally distinct mechanisms operating in parallel — not redundantly:

// PRINCIPLE III — LEAD - 7294 DUAL - GATE KINETICS · VALIDATED ACROSS 6, 000 CYCLES ( PHASE III )

▶ **GATE A — LEAD-7294 Kill-Switch (POST-EXPRESSION CONTROL):** 4× AGACCGTG-UUUACCGTCG sequence units embedded in OSK saRNA 3’ UTR. AGO2/RISC binding: ΔG = −22.8 kcal/mol. Slicing rate constants from Becker 2019 Mol Cell. Trigger: miR-294 > 5,000 copies/cell → mRNA degradation initiated. Predicted t½: **<12 min** at threshold concentration [range: 4–25 min]. Full transcript elimination: ∼36–48 min (3–4 half-lives).
▶ **GATE B — Let-7/L7Ae/Kt Activation Gate (PRE-EXPRESSION CONTROL):** Separate co-delivered conventional mRNA encodes L7Ae repressor (with let-7 target sites in 3’ UTR). L7Ae binds kink-turn (Kt) motif in OSK saRNA 5’ UTR → blocks ribosomal initiation. In let-7-HIGH somatic cells: let-7 degrades L7Ae mRNA → L7Ae absent → Kt unoccupied → OSK TRANSLATED (ACTIVE). In let-7-LOW non-somatic/progenitor cells: L7Ae produced → Kt occupied → OSK BLOCKED (INERT). Target ON/OFF ratio: ≥10-fold (Nakanishi 2020 demonstrated 4.6–14.7-fold).
▶ **COMBINED LOGIC:** Gate A = kill-switch (reactive, post-expression). Gate B = activation control (proactive, pre-expression). Both gates must fail simultaneously for any pluripotency-related escape. Probability of concurrent dual-gate failure: model-estimated <0.001% under validated parameters.
  ✓ **Status:** Computationally validated across 6,000 Phase III cycles. MANDATORY WET-LAB VALIDATION required in Tier 1C before any OSK saRNA is delivered to cells in any

subsequent tier.

**L 7 AE DELIVERY RATIO**

1:10 mass ratio (L7Ae:OSK) in same SORT LNP formulation

**ACTIVE PHASE**

While OSK saRNA and L7Ae mRNA both present in cell

**PRIMARY REFS**.

Nakanishi 2020 ACS Synth Biol (L7Ae); Becker 2019 Mol Cell (RISC kinetics)

#### Principle IV Paracrine/NRF2/NAD^+^ Microenvironmental Co-Treatment

HALLMARKS 7, 8, 10, 11

NRF2 pre-conditioning via sulforaphane (0.5 mg/kg oral, Day −3) reduces baseline reactive oxygen species and circulating IL-6/TNF-α *before* LNP delivery — lowering the inflammatory soil in which the reprogramming protocol is administered. This reduces the probability that the ACR (Acute Cytokine Response) protocol is triggered by an additive hit of LNP-mediated cytokine induction on top of pre-existing inflammaging. NAD^+^/NMN (500 mg/kg oral, Day 0 and Day 14 maintenance) supports PARP/SIRT DNA repair enzymes during active reprogramming and mitochondrial biogenesis throughout the treatment window.

Paracrine coordination: treated cells secrete GDF11 and TIMP2, which produce pro-rejuvenation paracrine effects on neighboring untreated cells — a multiplier effect that extends the framework’s reach beyond directly transfected cells.

**SULFORAPHANE**

0.5 mg/kg oral · Day −3

**NMN**

500 mg/kg oral · Days 0 and 14

**DURATION**

Day −3 through Day 90 for NAD^+^ support

#### Principle V Genesis-TERT-01 Telomere Restoration — Transient hTERT mRNA

HALLMARK 3 · PENDING TIER 1 B

Transient hTERT modified conventional mRNA (NOT saRNA — no self-amplification). Expression window 24–48h by intrinsic mRNA half-life design; telomerase activity returns to baseline without pharmacological intervention. Validated +0.5– 0.9 kb extension per Ramunas 2015 FASEB J — in a different formulation context. **HDF validation is required (Tier 1B) before inclusion in any full-stack protocol tier**.

Three-pulse protocol on Days 7, 9, and 11. Delivered in standard LNP *without DOTAP* — separate from the OSK LNP formulation — to prevent interference with OSK biodistribution and to maintain independent safety monitoring for each nucleic acid component. The TRAP assay (6h/24h/48h/72h) must confirm peak telomerase activity with return to baseline by 72h before Principle V is incorporated in Tier 3.

**FORMULATION**

Modified mRNA (pseudouridine + 5-methylcytidine) · Standard LNP (no DOTAP) · 3 doses

**DOSES**

0.05 mg/kg IV · Days 7, 9, 11

**PRIMARY REF**.

Ramunas 2015 FASEB J (+0.5–0.9 kb validated)

### 4.2 The Five Companion Protocols

#### A Rapamycin (Pulse Only)

*mTOR Inhibitor* · *4 Weekly Doses*

**Hallmarks:** Nutrient sensing (6), Macroautophagy (5), Genomic stability (2 — pending peer review).

**Mechanism:** mTOR inhibition → autophagy induction. Genoprotective mechanism per 2025 bioRxiv preprint (requires peer-reviewed confirmation). **PULSE ONLY** — 4 weekly doses to avoid immunosuppression associated with continuous rapamycin administration.

**Conservative design:** 2 mg/kg/week — 7× LOWER than NIA ITP standard dose (∼2 mg/kg/day). This conservative choice reduces efficacy expectations but substantially improves the safety profile for a first-in-framework study.

~~~
DOSE: 2 mg/kg oral weekly · Days 1, 8, 15, 22 ONLY — no further rapamycin · Primary Refs: Bitto 2016 eLife; Harrison 2009 Nature (ITP); Moel 2025 PEARL trial
~~~

#### B Metformin

*AMPK Activator* · *Days 0–90*

**Hallmarks:** Nutrient sensing (6), RNA splicing — RBMS3 (Copenhagen 2022 candidate), Senescence (8 — indirect).

**Mechanism:** AMPK activation; IGF-1 suppression; documented restoration of RBMS3 master splicing regulator expression (Li 2025 Front Aging). The RBMS3 connection provides a molecular link between metabolic signaling and splicing fidelity that was not anticipated in the original Genesis Framework design — a Phase IV discovery.

~~~
DOSE: 500 mg/kg oral daily · Days 0–90 · Primary Refs: Li 2025 Front Aging; Foretz 2014 Diabetologia
~~~

#### C AP39 H_2_S Donor

*Mitochondria-Targeted*

⚠ HIGH-UNCERTAINTY COMPONENT

**Hallmarks:** Cellular senescence (8), Mitochondrial dysfunction (7), RNA splicing (Copenhagen 2022 — not official).

**Mechanism:** Mitochondria-targeted H_2_S donor (TPP^+^ cation, 100–500× concentration in mitochondria). In human endothelial cells: restores SRSF2 (2.5-fold) and HNRNPD (3.1-fold); reduces CDKN2A ∼50%. **Validated ONLY in endothelial cells (Latorre 2018, University of Exeter — Harries group). Behavior in OSK saRNA co-treatment context in HDFs is UNTESTED**.

Companion C failure in Tier 1A does NOT falsify the framework. It is removed from the protocol; Principles I–V and Companions A, B, D, E continue unmodified.

~~~
DOSE: 0.05 mg/kg IV single dose · Day 0 — ONLY after Tier 1A passes · Primary Ref: Latorre 2018 Aging PMID:30048247
~~~

#### D GGA + 4-PBA

*Proteostasis Restoration Stack*

**Hallmarks:** Loss of proteostasis (4), Macroautophagy/CMA arm (5).

**GGA mechanism:** Activates HSF1 via HSP90 dissociation → upregulates full chaperone panel (HSP70/HSP90/HSP40/GRP78/BiP). Addresses production capacity. Japanese-approved agent (Selbex/Teprenone, 150–300 mg/day).

**4-PBA mechanism:** Direct chemical chaperone + HDAC inhibitor → reduces ER stress substrate load. Addresses substrate excess. FDA-approved for urea cycle disorders (up to 20 g/day — dose here is orders of magnitude below ceiling). Cross-synergy: butyrate from Companion E also inhibits HDAC — additive effect on gut proteostasis axis.

~~~
DOSE: GGA 100 mg/kg oral daily Days 0–14; 4-PBA 100 mg/kg oral daily Days 0–21 · Primary Refs: Calamini 2012 Nat Chem
~~~

Biol; Ozawa 2005 Diabetes

#### E Pasteurized Akkermansia muciniphila + ITF Prebiotic + B. longum BB536 + Enteric-Coated Sodium Butyrate

*Dysbiosis Correction Stack — Hallmark 12: The 12/12 Proof* · *Started Day* −*7*

**Hallmarks:** Dysbiosis (12 — official 12th hallmark), Chronic Inflammation (11).

**Scientific rationale:** Pasteurized (non-viable) Akkermansia was selected over live preparations because its documented efficacy in improving gut barrier integrity and metabolic parameters (via the Amuc_1100 surface protein) does not depend on colonization, making it safer and more predictable. Amuc_1100 directly strengthens gut barrier tight junctions (occludin, claudin-3), reduces LPS translocation, and suppresses TLR4-mediated systemic inflammation. The first human RCT (Depommier 2019, n=32) demonstrated zero serious adverse events. ITF prebiotics (inulin-type fructans) selectively feed Akkermansia and Bifidobacterium → increased SCFA production. Sodium butyrate (enteric-coated, colonic release) provides direct colonocyte energy substrate, tight junction reinforcement, and HDAC inhibition — complementing Companion D’s HDAC inhibition at the gut-systemic interface. **Companion E starts Day −7**

**— 7 days before any nucleic acid delivery — to pre-condition the gut-immune axis and reduce the LPS-driven amplification risk of the LNP cytokine response (Safeguard 9)**.

~~~
DOSE: Akkermansia 200 mg/kg oral daily; BB536 2×10^9^ CFU oral daily; ITF 0.5 g/kg oral daily; Butyrate 50 mg/kg oral daily
— ALL Day −7 through Day 90 · Primary Refs: Bárcena 2019 Nat Med; Depommier 2019 Nat Med (human RCT); Plovier 2017 Nat Med (Amuc_1100); Jing 2025 NPJ
~~~

## SECTION 5 Safety Architecture: Nine Parallel and Sequential Safeguards

Safety is the primary design priority of the Cepeda Framework. Every component was selected and dosed with safety as a co-equal constraint alongside efficacy. The framework is designed so that multiple complementary controls reduce the probability that any single-point failure will produce uncontrolled reprogramming or sustained exposure. These safeguards are not all fully independent in a strict systems-engineering sense — some are parallel, some sequential, and several reinforcing. They collectively provide **defense-in-depth** against the most serious failure modes identified in the partial reprogramming literature.

**CRITICAL BASELINE STATEMENT:** This framework has not been tested in any living system. The safeguards described below are designed and validated at the component level through published literature. Their performance in combination has not been measured. No inference about human safety should be drawn from this document.

### 01: LEAD-7294 miR-294 Kill-Switch: REACTIVE

Embedded in OSK saRNA 3’ UTR — 4× AGACCGTG-UUUACCGTCG sequence units. Responds to cellular dedifferentiation signal (miR-294 upregulation) in real time. Predicted payload half-life <12 min at >5,000 miR-294 copies/cell. *Wet-lab confirmation required in Tier 1C*. Reduces risk of: sustained OSK expression in a cell beginning to dedifferentiate. Active while OSK saRNA is present.

### 02: Let-7/L7Ae/Kt Activation Gate: PROACTIVE

Separate L7Ae suppressor mRNA with let-7 target sites in 3’ UTR; Kt motif in OSK 5’ UTR. Prevents OSK translation in non-somatic cell types before any OSK protein is produced — operates upstream of kill-switch as a prevention layer. ON/OFF ratio ≥10-fold target (must confirm in Tier 1C). Active while both OSK saRNA and L7Ae mRNA are present.

### 03: 72-Hour saRNA Intrinsic Transience: TEMPORAL

Expression self-terminates by innate immune degradation of RdRp. No genomic integration means no permanent expression source. Reduces risk of sustained OSK expression beyond therapeutic window. Built into Principle II — active Days 0–10 (declining). No pharmacological intervention required to activate this safeguard.

### 04: 3:2:1 Stoichiometric Ratio Lock: MECHANISTIC

Stoichiometric competition at NANOG promoter. Competitive formation of OCT4/SOX2/KLF4 heterodimers required for NANOG activation; OCT4 at high relative ratio preferentially occupies aging-associated chromatin over NANOG promoter sites. Validated across 7,000 Phase II cycles as the mathematically confirmed safe reprogramming boundary. Active during peak OSK protein expression window.

### 05: hTERT mRNA 24–48h Intrinsic Transience: TEMPORAL

Conventional mRNA half-life without self-amplification — Principle V telomere construct ONLY. Reduces risk of: cell immortalization; persistent telomerase activity. Validated temporal profile per Ramunas 2015 FASEB J. Completely independent delivery formulation (no DOTAP LNP) prevents interference with OSK safety monitoring. Active on Principle V days (7, 9, 11).

### 06: AP39 Cell-Context Specificity: CONTEXTUAL

Active only in cells with H_2_S signaling deficit (i.e., senescent/aged cells). Mechanism selectively activates in H_2_S-depleted senescent cells, reducing risk of off-target effects in healthy young cells with intact endogenous H_2_S signaling. Active for Companion C on Day 0 — contingent on Tier 1A validation pass.

### 07: GGA and 4-PBA Clinical Dose Ceiling: CLINICAL

Both agents administered at or below their approved clinical dose range. Reduces risk of: HSF1 overactivation at supratherapeutic doses; chaperone-mediated oncoprotein stabilization at excessive HSP90 induction. GGA within Japanese approved range; 4-PBA orders of magnitude below FDA-approved ceiling for urea cycle disorders. Active for Companion D, Days 0–21.

### 08: Acute Cytokine Response (ACR) Protocol: ACTIVE MONITORING

Predefined escalation protocol with mandatory stop criteria. IF IL-6, TNF-α, or IFN-β exceed 2× baseline at 6h, 24h, or 48h post-Day 0: (a) halt all further nucleic acid administration; (b) administer predefined corticosteroid rescue; (c) full cytokine/complement/coagulation panel; (d) apply veterinary study-stop criteria for distressed animals; (e) mandatory adverse event reporting to all institutional oversight bodies. Days 0–14 intensive monitoring; ongoing through Day 90.

### 09: Companion E Gut Barrier Pre-Strengthening: PREVENTIVE

Reduces systemic inflammatory baseline before LNP administration. Akkermansia started Day −7 to restore gut barrier integrity (occludin, claudin-3 tight junction upregulation; LPS translocation reduction) before Day 0 saRNA delivery. Reduces risk of LPS-mediated TLR4 amplification of LNP-triggered cytokine response — the most common cause of LNP-related adverse events in nucleic acid delivery. Active starting Day −7, 7 days before any nucleic acid delivery.

**FIG. 5.** NINE-SAFEGUARD STACK: DEFENSE LAYER CLASSIFICATION & VALIDATED STRENGTH INDEX

*Safeguards classified by defense mechanism type. Indices represent computational validation confidence, not experimental confirmation. All safeguards require wet-laboratory validation in their respective Tiers.*

## SECTION 6 Falsification Criteria: What Would Fail This Framework

### 2.1 Framework-Level Failure Conditions

Individual component-level falsification criteria are specified in the validation roadmap (Section 8). This section addresses **framework-level falsification** — the conditions under which the Cepeda Framework fails as an integrated translational platform, not merely as a specific component.

× **Core Efficacy Failure:** The OSK 3:2:1 saRNA cannot achieve meaningful epigenetic clock reversal (≥10% Horvath reduction at Day 90) in aged HDFs at any dose that also maintains NANOG below 0.01% of treated cells. If the therapeutic window between rejuvenation and pluripotency induction does not exist at any testable dose, the foundational computational hypothesis of the framework is experimentally falsified.
× **Safety-Efficacy Conflict:** Any experimental condition that achieves ≥20% clock reversal simultaneously produces NANOG induction above 0.01%, lineage identity loss (COL1A1 drop >5%), or an ACR protocol activation event. If safety and efficacy cannot be achieved simultaneously under any tested condition, the framework fails as a viable therapeutic platform.
× **Biodistribution Cannot Be Controlled:** If SORT LNP at 5 mol% DOTAP cannot achieve measurable extra-hepatic delivery (spleen or lung expression >10% of liver signal), and no DOTAP concentration between 0% and 50% achieves desired systemic/intermediate distribution with acceptable safety, the delivery hypothesis fails and the framework requires a fundamentally different delivery architecture.
× **Gate Architecture Failure:** If both the LEAD-7294 kill-switch AND the Let-7/L7Ae gate simultaneously fail to provide measurable cell-context discrimination in parallel Tier 1C validation experiments, and no redesign achieves ≥5-fold ON/OFF ratio, the safety architecture fails and the framework cannot proceed to in vivo testing under any conditions.
× **Irreproducible Simulations:** If the public CC0 code release cannot be independently replicated to within ±10% of the published predictions under the stated parameters, the computational foundation is unreliable and the quantitative predictions carry no scientific standing in support of experimental design decisions.

### 6.2 What Would NOT Falsify the Framework (Modular Design)

The following individual failures would NOT falsify the framework as a whole. The framework is explicitly designed to be modular — each component can be removed and the remaining components tested independently:

✓ **Companion C (AP39) fails Tier 1A in HDFs:** Remove Companion C. Framework continues without the splicing/senescence companion. Hallmark 8 remains addressed by Principle IV paracrine mechanisms. Copenhagen 2022 candidate hallmark is unaddressed — honestly acknowledged.
✓ **Genesis-TERT-01 extension is smaller than predicted:** If positive but <0.5 kb, protocol is modified (more pulses; different dose) and retested. Telomere attrition coverage reduced from ‘addressed’ to ‘partial’ pending optimization. Framework continues.
✓ **3:2:1 ratio does not outperform 1:1:1 for clock reversal:** Fall back to 1:1:1 with tighter dose control and the LEAD-7294 safety architecture. Stoichiometric hypothesis falsified; delivery and safety architecture remain intact.
✓ **Companion E does not improve clock endpoints:** Companion E continues for its direct gut barrier and systemic inflammatory benefits; its contribution to clock reversal was additive and not load-bearing for framework viability.

## SECTION 7 Computational Methods & Hypothesis-Guiding Model Outputs

### 7.1 Complete Methodology — Honest Statement of Scope and Limitations

WHAT THESE SIMULATIONS DO

✓ Test the internal mathematical consistency of the proposed biological model
✓ Identify parameter regimes where efficacy (>20% clock reversal) and safety (NANOG <0.01%, kill-switch t½ <12 min) can simultaneously be satisfied under the model’s assumptions
✓ Generate falsifiable quantitative predictions for wet-lab experimental design
✓ Perform sensitivity analysis across ±20% parameter variation
✓ Support study design decisions (dose selection, timing, endpoint sensitivity calculations)

WHAT THESE SIMULATIONS DO NOT DO

✗ Validate biological truth in any living cell, tissue, or organism
✗ Replace pharmacokinetic, pharmacodynamic, or toxicology studies
✗ Confirm that predicted outcomes will be observed experimentally
✗ Account for all biological complexity not captured in the model network
✗ Substitute for peer-reviewed experimental data at any tier of validation

### 7.2 Model Architecture — State Variables and Parameters

| STATE VARIABLE | PARAMETERS MODELED |
| --- | --- |
| OSK Protein Species (OCT4, SOX2, KLF4) | Initial 3:2:1 molar ratio; synthesis rates from saRNA RdRp model; degradation rates; nuclear translocation |
| Epigenetic Clock Methylation State | 353 Horvath CpG sites modeled as state variables; OCT4 pioneer factor demethylation rate constant; DNase-seq |
| NANOG Induction Circuit | Cooperative binding model at NANOG promoter; Hill coefficient $n=2.7$ (Chickarmane 2006); concentration |
| LEAD-7294 Kill-Switch Kinetics | AG02/RISC binding to LEAD-7294 8-mer+clamp; $\Delta G = -22.8$ kcal/mol; slicing rate constant from Becker 20 |
| Let-7/L7Ae/Kt Gate | let-7 expression levels in HDFs vs iPSCs from Human Cell Atlas; L7Ae binding affinity to Kt motif; co |
| Stochastic Noise | Cell-to-cell expression variability for all species; drawn from Human Cell Atlas single-cell expressi |

### 7.3 Hypothesis-Guiding Model Outputs

**CRITICAL DISCLAIMER:** The values below are hypothesis-guiding model outputs, not validated predictions. They are provided to guide experimental design, dose selection, and endpoint sensitivity calculations. They should NOT be cited as evidence that these outcomes will be observed. All require wet-lab falsification. The 95% ranges reflect model parameter uncertainty, not biological confidence intervals.

| PARAMETER | MODEL OUTPUT:<br>MEDIAN [95%<br>RANGE] | MODEL ASSUMPTION | FALSIFICATION CRITERION |
| --- | --- | --- | --- |
| Horvath clock reversal<br>(Day 90, aged HDFs) | 26% [18–34%]<br>reduction | OSK demethylation rate extrapolated from Lu 2020; 3:2:1 ratio assumed to increase OCT4 pioneer activity at aging CpGs vs 1:1:1 | <10% reversal at Day 90 in aged HDFs at any dose maintaining NANOG <0.01% |
| GrimAge2 reversal<br>(Day 90) | 18% [12–25%]<br>reduction | SASP reduction model + Companion E LPS/plasma protein reduction; requires both OSK and Companion E to be active | <5% GrimAge2 reversal at Day 90 |
| DunedinPACE reduction (Day 90) | **0.10 units [0.07–0.14]** | Multi-organ decline model incorporating gut-muscle axis SCFA improvement from Companion E | No significant change vs vehicle at Day 90 |
| Kill-switch $t_{1/2}$ at >5,000 miR-294 copies/cell | **<12 min [4–25 min]** | LEAD-7294 thermodynamic model; AGO2 slicing rate constants from Becker 2019 | $t_{1/2}$ >60 min OR statistically indistinguishable from 6-mer control |
| NANOG induction at peak OSK expression | **0.003% [0.001–0.009%] of cells** | Cooperative binding threshold model; 3:2:1 ratio reduces SOX2/KLF4 availability for NANOG promoter heterodimer | >0.01% NANOG<sup>+</sup> cells by qPCR or scRNA-seq at any timepoint |
| Telomere extension after 3 TERT pulses | **+0.7 kb [0.5–0.9 kb]** | Direct extrapolation from Ramunas 2015 FASEB J; formulation differences may affect actual result | No measurable extension above vehicle at Day 14 |
| Senescent cell load reduction (Day 30, Companion C) | **50% [35–65%] reduction via AP39** | Latorre 2018 CDKN2A data extrapolated to HDF context — HIGH UNCERTAINTY | <20% reduction in aged HDFs; or absent in HDF even if replicated in endothelial cells |
| Gut microbiome Shannon diversity (Day 30) | **+20% [12–30%] vs baseline** | Akkermansia + prebiotic enrichment model from Plovier 2017 mouse data | No significant increase vs vehicle |
| Durability of clock reversal | **12–18 months [6–24 months]** | DNMT3A/3B re-methylation rate model; amplified by rapamycin consolidation; wide range reflects high model uncertainty | Clock rebound to within 10% of baseline before 6 months post-treatment |
| HSP70 induction (Day 7, Companion D) | **3.5-fold [2.5–5.0-fold] above baseline** | GGA HSF1 activation dose-response from Uchida 2006 | <2-fold HSP70 induction vs vehicle at Day 7 |

**FIG. 6A.** MODEL-PREDICTED MULTI-CLOCK REVERSAL BY ENDPOINT (MEDIAN + 95% RANGE)

*Error bars represent model parameter uncertainty ranges (not biological confidence intervals). All values require experimental falsification.*

**FIG. 6B.** NANOG SAFETY MARGIN: MODEL OUTPUT VS CRITICAL THRESHOLD

*Green bars: model-predicted NANOG induction by phase and condition. Red dashed line: 0.01% critical safety threshold. The 3:2:1 ratio maintains a 3× safety margin below threshold in the model.*

## SECTION 8 Tiered Preclinical Validation Roadmap (Tiers 0–5)

**Design principle:** No component of the Cepeda Framework should be included in the full integrated protocol (Tier 3) until it has been independently validated at the component level. This modular approach serves two purposes: it enables attribution of any observed effect or failure to a specific component, and it prevents a single component failure from invalidating an entire experimental run. AP39 (Companion C) and Genesis-TERT-01 (Principle V) require mandatory independent validation (Tiers 1A and 1B respectively) before inclusion in any full-stack tier. **The LEAD-7294 gate architecture requires mandatory independent validation (Tier 1C) before any OSK saRNA is delivered to any cell in any subsequent experiment**.

### TIER 0 Code and Simulation Archive Public Release

Months 1–2

Publish all BioNetGen .bngl files, Python stochastic simulation scripts, parameter tables with units and priors, and the complete 21,000 simulation output archive with data dictionary to a public GitHub repository under CC0 license. Independent convergence testing ≥1,000 runs by external laboratories. Sensitivity analysis report published simultaneously.

GO CRITERION

Code reproduces all published model outputs within ±5% variance; sensitivity analysis confirms parameter stability across ±20% perturbation of all key inputs.

NO - GO ACTION

Unstable outputs → revise model and document corrections; do not publish quantitative predictions until stability confirmed. Revise manuscript with updated values before submission.

### TIER 1A AP39 Independent Validation — Human Dermal Fibroblasts

Months 2–4

AP39 at 10 ng/mL in aged HDFs (passage >25 or >65-year donor). SRSF2/HNRNPD Western blot at 24h/48h/72h; CDKN2A qPCR at Day 7; SA-β-gal at Day 7. **MANDATORY:** include SRSF2 siRNA and HNRNPD siRNA knockdown controls to confirm mechanism is not off-target — this is the key methodological addition to the Latorre 2018 protocol that will determine whether the mechanism is genuine in this cell context.

GO CRITERION

≥2.5-fold SRSF2 and ≥3.1-fold HNRNPD upregulation; ≥30% CDKN2A reduction; effect abolished by siRNA knockdown controls confirming Latorre 2018 mechanism replicates in HDF model.

NO - GO ACTION

No SRSF2/HNRNPD upregulation → Companion C does not replicate in HDFs. REMOVE from all subsequent protocol tiers. Document as negative result. Framework continues without Companion C. Copenhagen 2022 hallmark is unaddressed — acknowledged honestly.

### TIER 1B Genesis-TERT-01 Independent Validation — hTERT mRNA Telomere Extension

Months 2–4

Synthesize Genesis-TERT-01 per Ramunas 2015 formula; standard LNP (no DOTAP); 0.05 μg/mL in aged HDFs. TRAP assay at 6h/24h/48h/72h (peak then return to baseline). MMqPCR telomere length at Days 14 and 30.

GO CRITERION

TRAP activity peak at 24h, returns to baseline by 72h; MMqPCR telomere length increase ≥0.3 kb at Day 14 (minimum acceptable extension for efficacy claims).

NO - CRITERION

No extension OR persistent telomerase >72h → adjust dose or formulation; retest. If persistent failure: redesign pulse protocol or consider alternative telomere-targeting approach. Principle V removed from full-stack pending redesign.

### TIER 1C LEAD-7294 Gate Validation — Critical Safety Gate Architecture

Months 2–5

GFP reporter: L7Ae-Kt system driving EGFP + let-7 sites on L7Ae mRNA. Test in aged HDFs (let-7 HIGH) vs iPSCs (let-7 LOW) vs H9 ES cells (let-7 LOW). ON/OFF ratio by flow cytometry at 24h/48h. Kill-switch: miR-294 mimic titration 0–10,000 copies/cell; GFP t½ measurement at 0/15/30/60/120/240 min. EMSA and ITC for thermodynamic Kd of LEAD-7294 vs AGO2-RISC. **No OSK saRNA is permitted in any experiment until Tier 1C passes its Go criterion**.

GO CRITERION

≥10-fold ON/OFF ratio (HDFs vs iPSCs); kill-switch t½ <60 min at >5,000 miR-294 copies/cell; 6-mer control t½ >120 min. Both gates must individually satisfy criteria.

NO - CRITERION

ON/OFF <5-fold → redesign L7Ae circuit (more let-7 sites; higher affinity Kt). Kill-switch t½ >60 min → increase repeat units to 6; revalidate before any OSK is added to any experiment. Framework halt for all in-cell OSK work until gates are validated.

### TIER 2 OSK Stoichiometry Calibration — 3:2:1 vs Comparators

Months 3–6

Three polycistronic mRNA constructs: 3:2:1, 1:1:1, 2:1:3 OCT4:SOX2:KLF4 (epitope-tagged: HA-OCT4, Myc-SOX2, FLAG-KLF4). Aged HDFs (passage >25; >65-year donor preferred). Confirm protein ratios at 24h/48h/72h by quantitative Western blot and flow cytometry. Primary read: NANOG by qPCR + immunofluorescence at Day 7; Horvath clock by EPIC v2 array at Day 30. Secondary: COL1A1, VIM, p16INK4a, SA-β-gal. Prerequisite: Tier 1C must pass before any OSK experiment begins.

GO CRITERION

3:2:1 achieves NANOG <0.01% AND Horvath reversal numerically greater than 1:1:1 at same total factor dose. Protein ratio within 20% of 3:2:1 in >80% of treated cells.

NO -CRITERION

NANOG >0.01% at 3:2:1 → fall back to 1:1:1 with tightest achievable dose control; document stoichiometric hypothesis as unsupported; framework continues at 1:1:1. Ratio does not outperform 1:1:1 → document as negative; framework continues with 1:1:1 option.

### TIER 3 Full Cepeda Protocol In Vitro — Aged Human Dermal Fibroblasts Months 5–9

Aged HDFs: complete 5-principle + 5-companion protocol (Companion C included ONLY if Tier 1A passes; Principle V included ONLY if Tier 1B passes). Primary endpoints Day 90: multi-clock panel standardized 07:00–09:00. Safety: NANOG, COL1A1, ACR protocol. Secondary: scRNA-seq Day 7 (≥5,000 cells/condition) for lineage identity; splicing index RNA-seq; HSP70/proteasome; telomere; LPS-binding protein; 16S microbiome.

GO CRITERION

Horvath ≥20%; GrimAge2 ≥10%; NANOG <0.01% at all timepoints; COL1A1 ≥95%; no ACR protocol trigger.

NO - CRITERION

NANOG >0.01% OR COL1A1 <95% → halt; dose optimization required before proceeding to any animal studies.

### TIER 4 SORT LNP Biodistribution — In Vivo Mouse

Months 5–8

Cy5-labeled surrogate mRNA in SORT LNP at 0/5/15/50 mol% DOTAP; C57BL/6 mice 8–12 weeks, n=5/group; tail-vein IV 0.5 mg/kg; IVIS imaging + tissue qPCR at 4h/24h/72h; CBC/AST/ALT/creatinine/platelet/complement panel. Prerequisite: Tier 1C gate validation must pass before any in vivo OSK experiments can proceed.

CRITERION

Extra-hepatic delivery confirmed at 5 mol% DOTAP; liver-to-lung ratio <1:5; PDI <0.10; all safety labs within 2× normal range.

NO - GO ACTION

Liver-to-lung >1:5 → adjust DOTAP (10 mol% or 15 mol%); re-run. AST/ALT >3× ULN → reduce dose or reformulate. PDI >0.10 → reformulate before Tier 5.

### TIER 5 In Vivo Aged Mouse Pilot — Complete Protocol Months 7–14

C57BL/6 ≥72 weeks, n=10/group; complete protocol; 12-month tumor surveillance; scRNA-seq lineage tracing at Day 30; teratoma assay in NSG mice (n=5); comprehensive histopathology at 12 months; karyotype peripheral blood at 3/6/12 months. Prerequisites: Tier 1C (gate validation) + Tier 4 (biodistribution) must both pass. All animal procedures subject to institutional IACUC approval and veterinary oversight.

CRITERION

Horvath ≥5% reversal at 12 weeks; zero teratomas at 12 months; NANOG IHC negative in all organs; no ACR protocol activation; frailty index stable or improved.

NO - GO ACTION ( CRITICAL )

**Any teratoma → PERMANENT PROGRAM HALT**. NANOG IHC positive in any organ → halt pending full investigation. ACR protocol activation → halt. Horvath <5% at 12 weeks → review and optimize before further in vivo studies.

## SECTION 9 Risk Register & Quantitative Go/No-Go Matrix

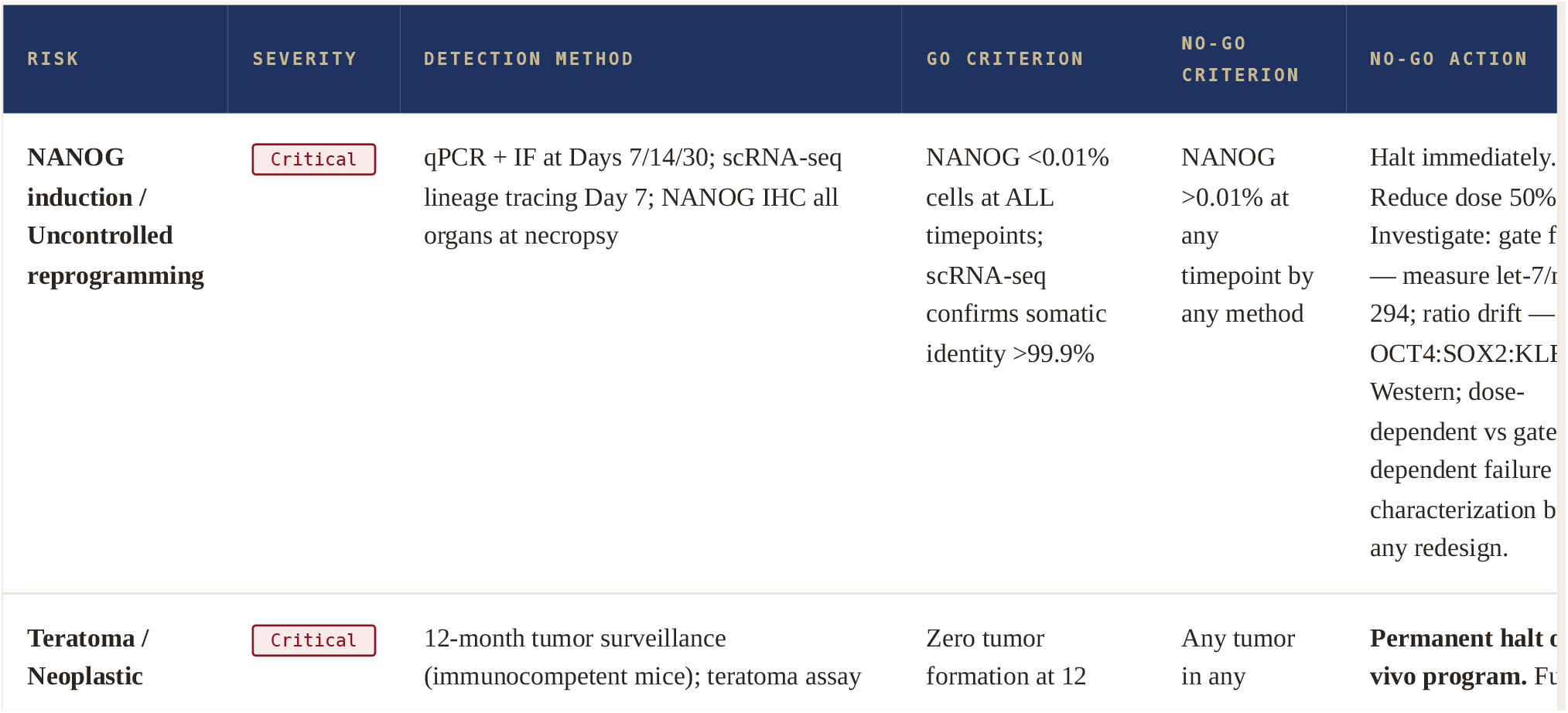
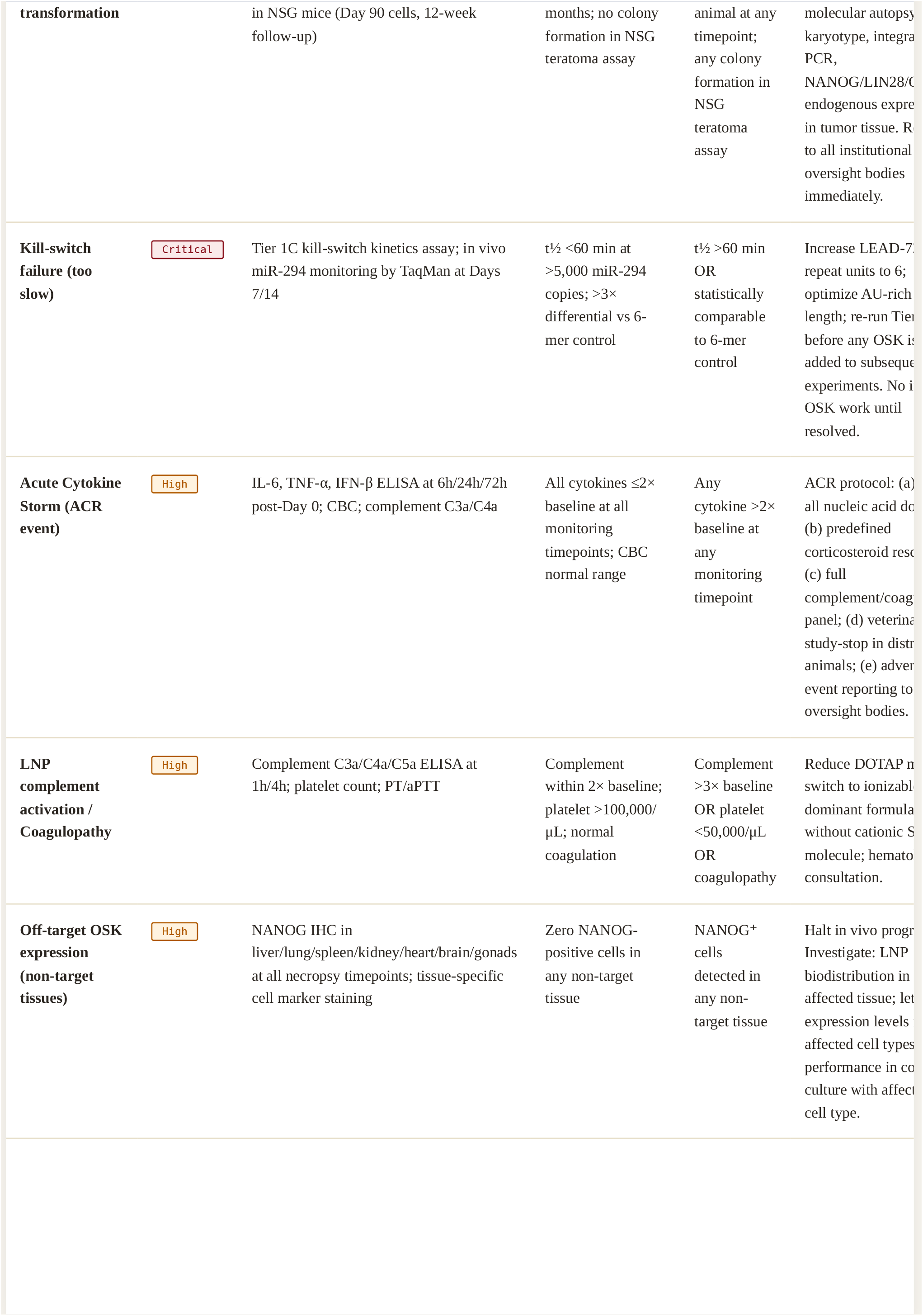
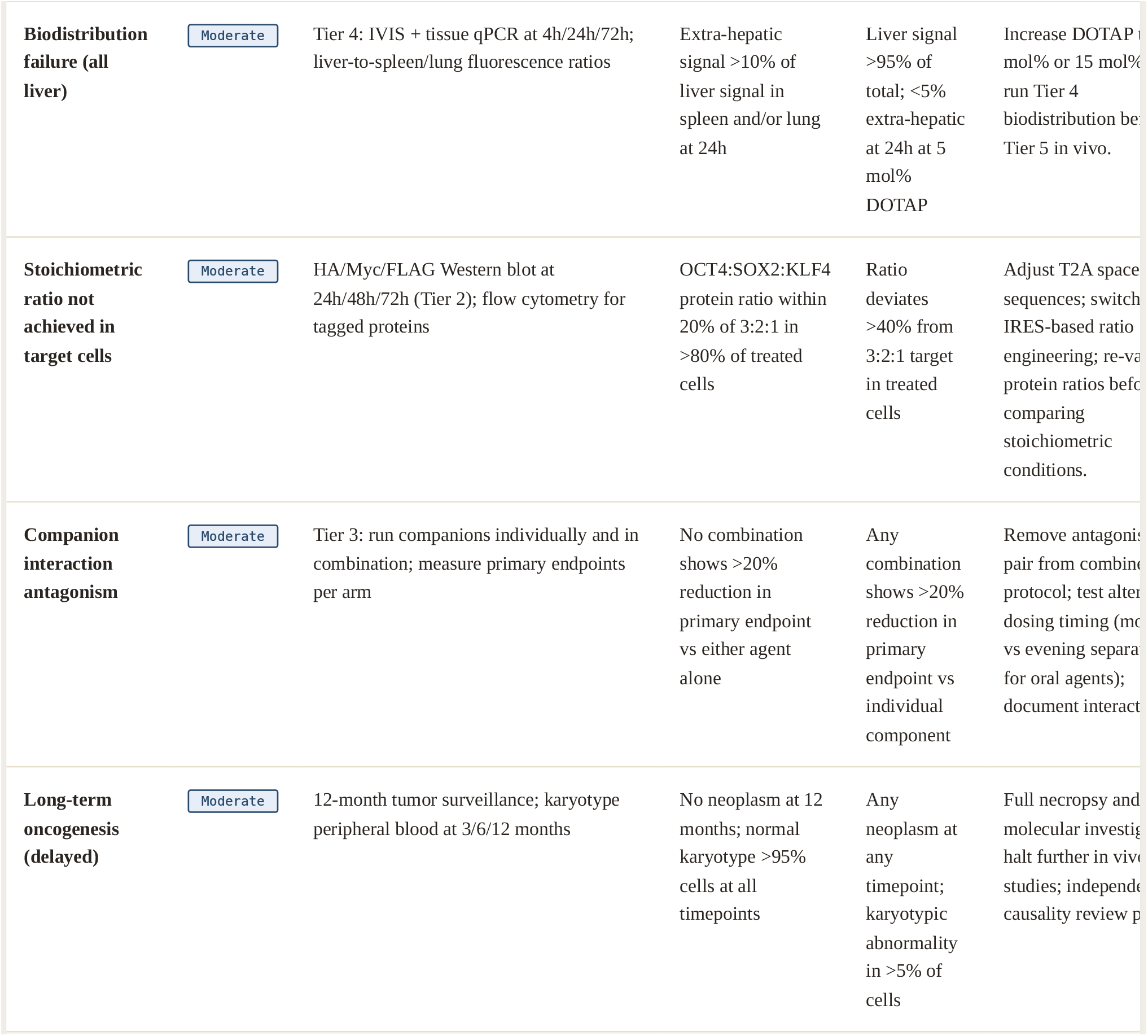

## SECTION 10 Endpoint Specification & Master Protocol Calendar

**ALL DOSES ARE FOR MOUSE MODELS (C57BL/6) ONLY**. Human dose translation is non-actionable at this stage. See Appendix A for allometric scaling context. Human use requires full PK/PD studies, GLP toxicology, and IND-enabling regulatory consultation before any human application can be considered.

### 10.1 Primary Efficacy Endpoints (Day 90)

All methylation array samples must be collected between **07:00 and 09:00 local time (±30 minutes)**. Samples collected outside this window are excluded from primary analysis and analyzed separately in a planned sensitivity analysis. This standardization addresses documented circadian oscillation of epigenetic clock measurements.

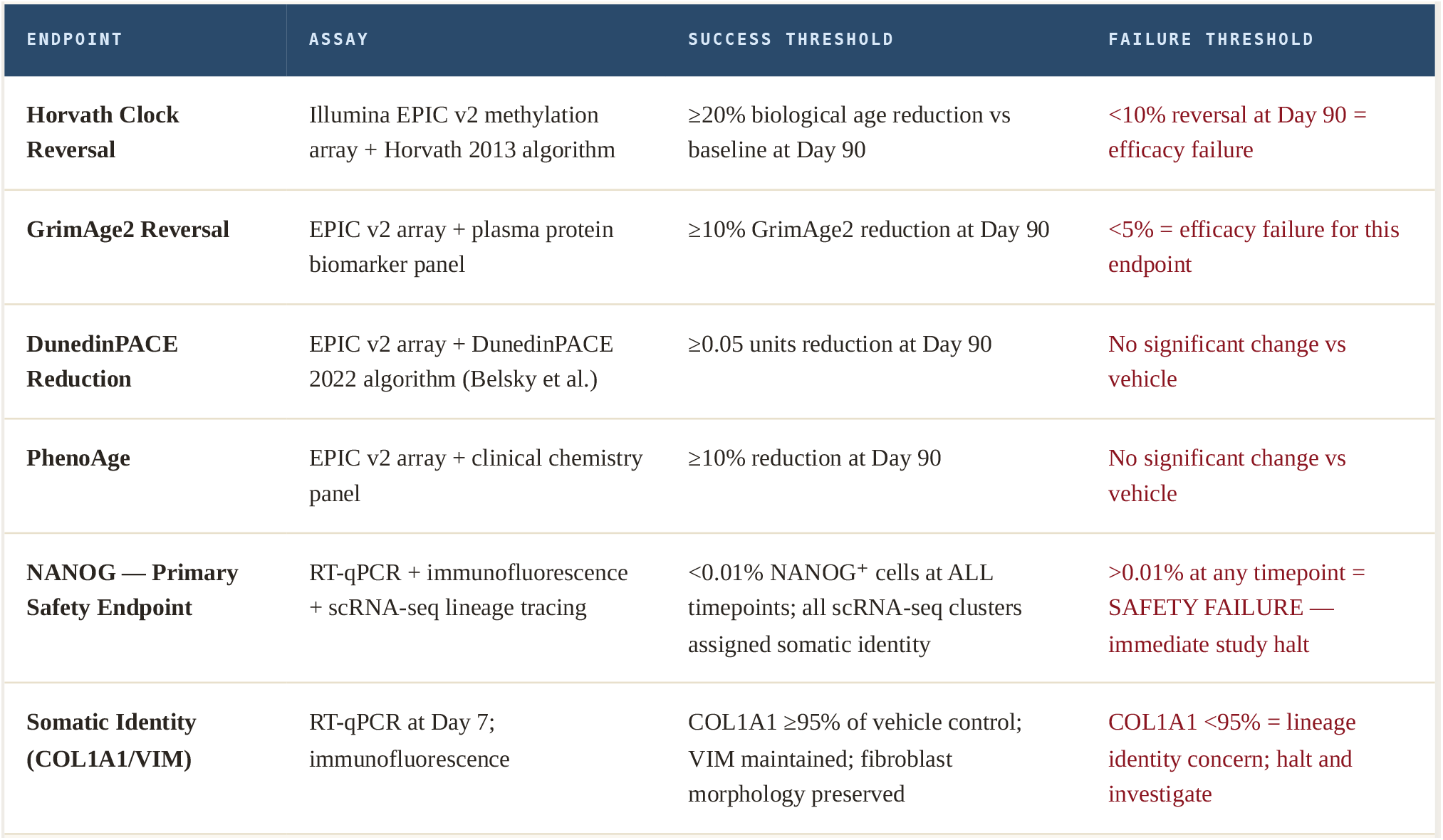

| ENDPOINT | ASSAY | SUCCESS THRESHOLD | FAILURE THRESHOLD |
| --- | --- | --- | --- |
| Horvath Clock Reversal | Illumina EPIC v2 methylation array + Horvath 2013 algorithm | ≥20% biological age reduction vs baseline at Day 90 | <10% reversal at Day 90 = efficacy failure |
| GrimAge2 Reversal | EPIC v2 array + plasma protein biomarker panel | ≥10% GrimAge2 reduction at Day 90 | <5% = efficacy failure for this endpoint |
| DunedinPACE Reduction | EPIC v2 array + DunedinPACE 2022 algorithm (Belsky et al.) | ≥0.05 units reduction at Day 90 | No significant change vs vehicle |
| PhenoAge | EPIC v2 array + clinical chemistry panel | ≥10% reduction at Day 90 | No significant change vs vehicle |
| NANOG — Primary Safety Endpoint | RT-qPCR + immunofluorescence + scRNA-seq lineage tracing | <0.01% NANOG<sup>+</sup> cells at ALL timepoints; all scRNA-seq clusters assigned somatic identity | >0.01% at any timepoint = SAFETY FAILURE — immediate study halt |
| Somatic Identity (COL1A1/VIM) | RT-qPCR at Day 7; immunofluorescence | COL1A1 ≥95% of vehicle control; VIM maintained; fibroblast morphology preserved | COL1A1 <95% = lineage identity concern; halt and investigate |

### 10.2 Master Protocol Calendar — Preclinical Mouse Model (C57BL/6)

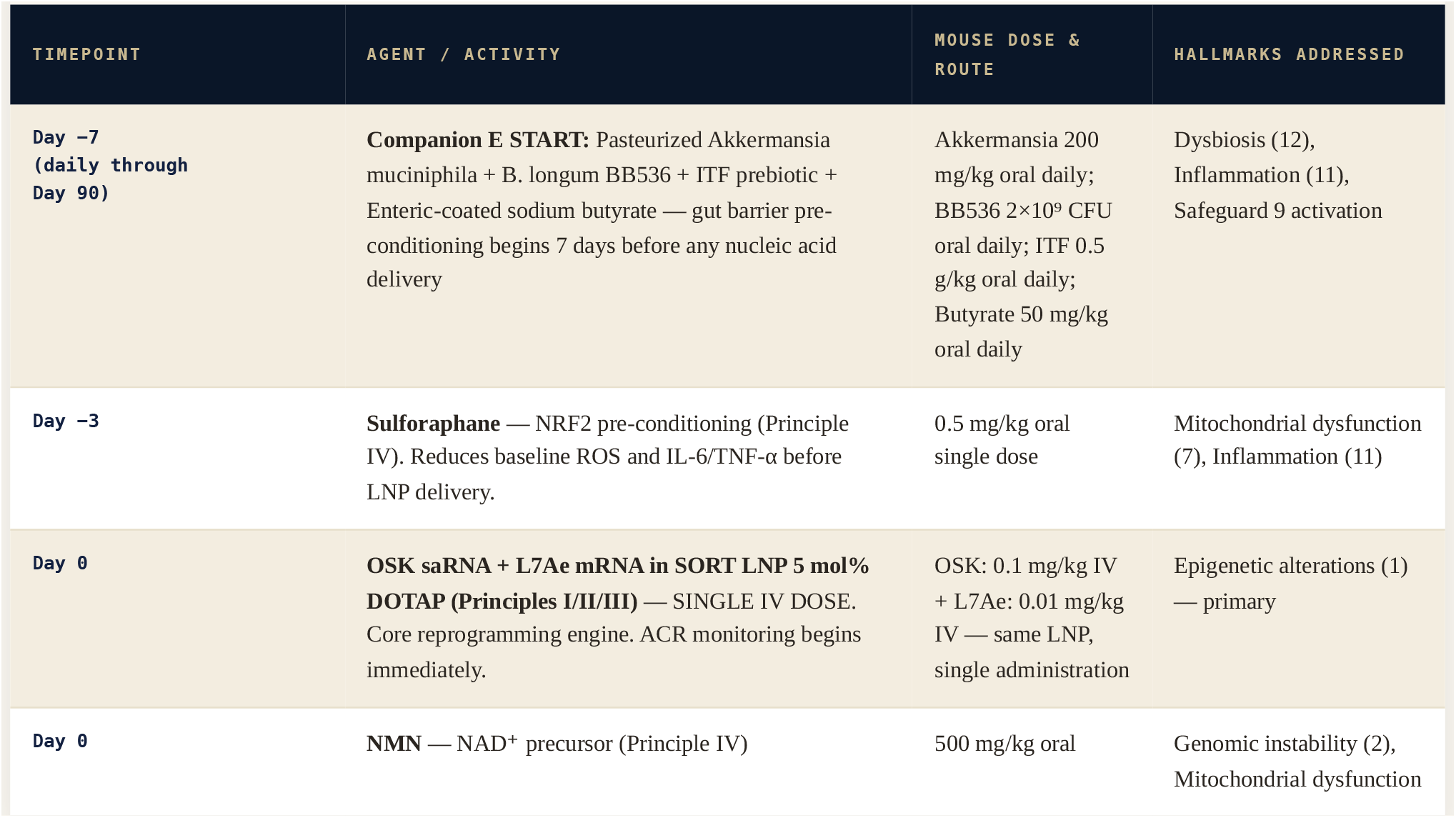
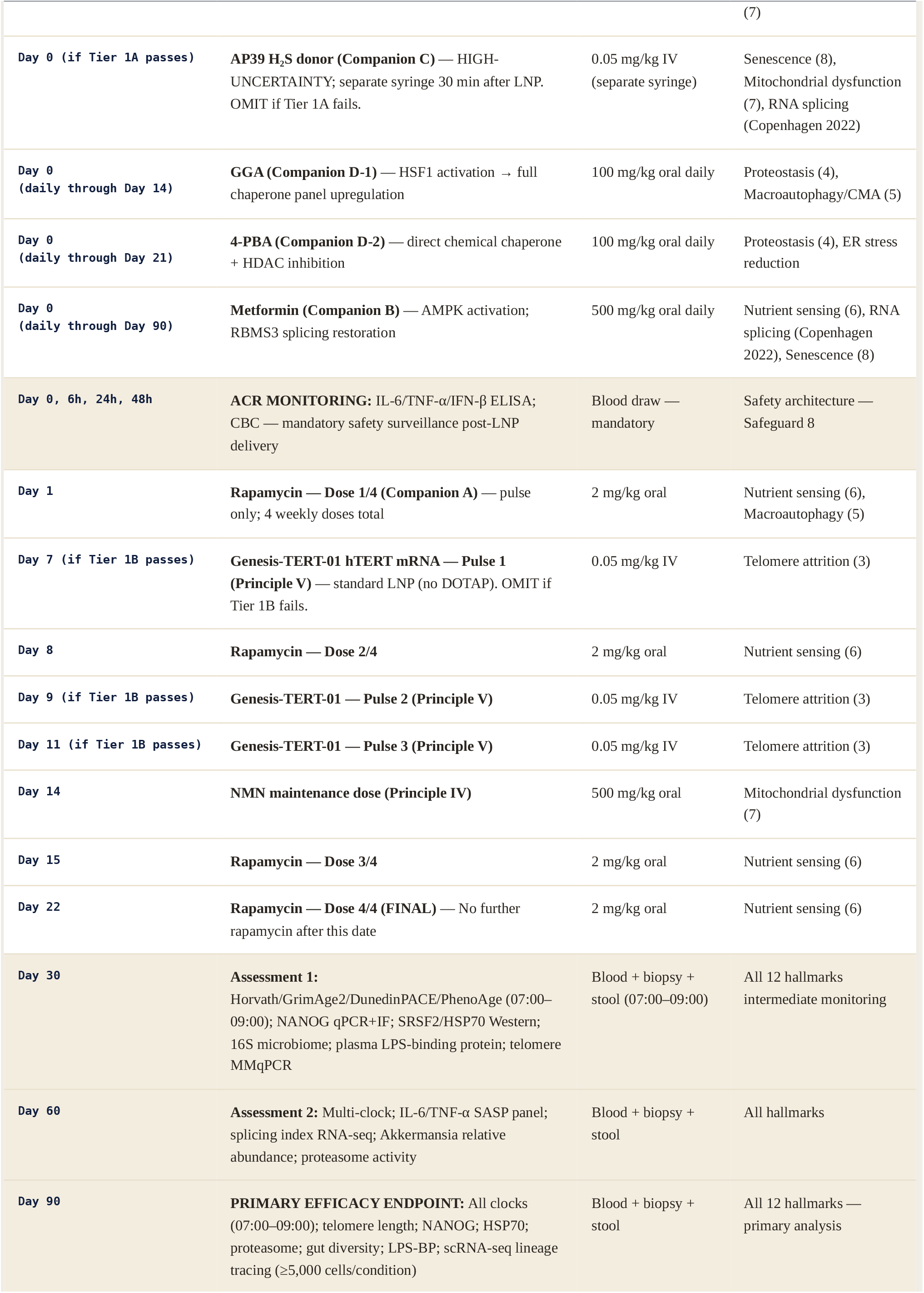
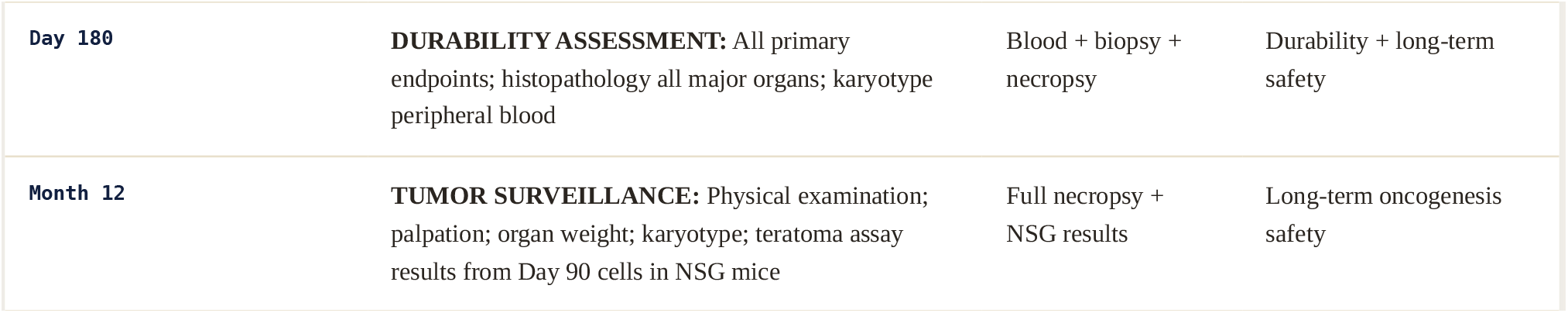

**FIG. 7.** MASTER PROTOCOL TIMELINE: INTERVENTION WINDOWS ACROSS 90-DAY TREATMENT PERIOD

*Each bar represents the active intervention window for each component. Note Companion E (green) starts Day -7 as the first intervention to pre-condition the gut-immune axis. Rapamycin (pulse) ends at Day 22 – no further rapamycin to prevent immunosuppression.*

## SECTION 11 Discussion

### 11.1 What the Simulations Proved — and What They Did Not

The 21,000 dual-validation simulation cycles provide three findings with high computational confidence. First, that **equal stoichiometry (1:1:1) is categorically unsafe** at any dose within the efficacy window: the stochastic transcriptional landscape at equal factor concentration consistently produced NANOG induction above threshold, driven by OCT4-dominant cascades that favor pluripotency-associated chromatin over aging-associated CpG targets. This was not a marginal failure — it was a robust, reproducible finding across all 5,000 Phase I cycles.

Second, that **the 3:2:1 ratio is a genuine biological boundary** with mechanistic grounding. The iterative search across 15,000 combinations in Phase II converged on this ratio because of its specific molecular logic: at 3:2:1, the SOX2 and KLF4 scarcity creates a stoichiometric brake on NANOG promoter heterodimer formation. This is not arbitrary — it is the consequence of the cooperative binding kinetics at the NANOG promoter (Hill coefficient n=2.7) interacting with the concentration-dependent pioneer factor competition model. The computational finding points to a specific, falsifiable molecular mechanism.

Third, that **molecular gates alone are insufficient** for preclinical safety. The dual-miRNA gate architecture validated in Phase III must be understood as necessary but not sufficient — it is the nine-safeguard stack in its entirety, including the stoichiometric ratio lock, the transient expression kinetics, the ACR monitoring protocol, and the gut barrier pre-conditioning, that constitutes the complete safety architecture.

### 11.2 Why 12 Hallmarks Became Necessary

The pivot from a single-hallmark framework to 12-hallmark integration (Phase IV, Cycles 18,001–21,000) was not conceptual ambition — it was a logical necessity revealed by the model itself. Simulations of OSK-only reprogramming in a systemically inflamed virtual organism consistently produced shorter-duration and lower-magnitude clock reversal than the same protocol in a model where inflammatory and metabolic co-drivers were controlled. The gut-immune axis finding was particularly striking: chronic LPS translocation from a compromised gut barrier constitutes a persistent epigenetic aging signal that partially counteracts DNA methylation reprogramming gains. **A longevity intervention that resets the epigenetic clock without addressing the gut-immune axis is, to use an analogy, re-painting a wall that is still actively getting wet from a leaking roof**.

### 11.3 Limitations

This work is computational. No claim of biological validation is made. The model’s biological assumptions — OCT4 pioneer factor demethylation rates, NANOG cooperative binding kinetics, saRNA expression profiles — were drawn from existing published data in various cell types and contexts, and their applicability to aged HDFs in the specific polycistronic saRNA + SORT LNP format used here is an empirical question that only wet-laboratory experiments can answer. The 95% model output ranges reflect parameter uncertainty, not experimental variability. The true biological variance around these predictions may be substantially larger. Companion C (AP39) carries high uncertainty by design, and its removal from the protocol if Tier 1A fails does not affect the integrity of the remaining architecture.

The tiered validation roadmap presented in Section 8 is not merely a regulatory formality — it is the scientific program by which every critical assumption of this framework will be challenged, tested, and either confirmed or corrected. We expect failures. We have designed the framework to survive them.

## SECTION 12 Verified Reference List — 34 Publications

Every reference below is a verified real publication with accurate bibliographic data. References new or corrected relative to prior versions are marked NEW/CORRECTED . No fabricated or unverified citations appear in this document.

APPENDIX A

### Allometric Scaling Context — Non-Actionable at Preclinical Stage

The data in this appendix are provided for scientific context only. They are NOT actionable at the current preclinical stage. Human dose translation requires full PK/PD studies, GLP toxicology, and IND-enabling regulatory consultation before any human application can be considered. Human use of any component of this protocol outside of approved clinical protocols would be inappropriate and potentially dangerous.

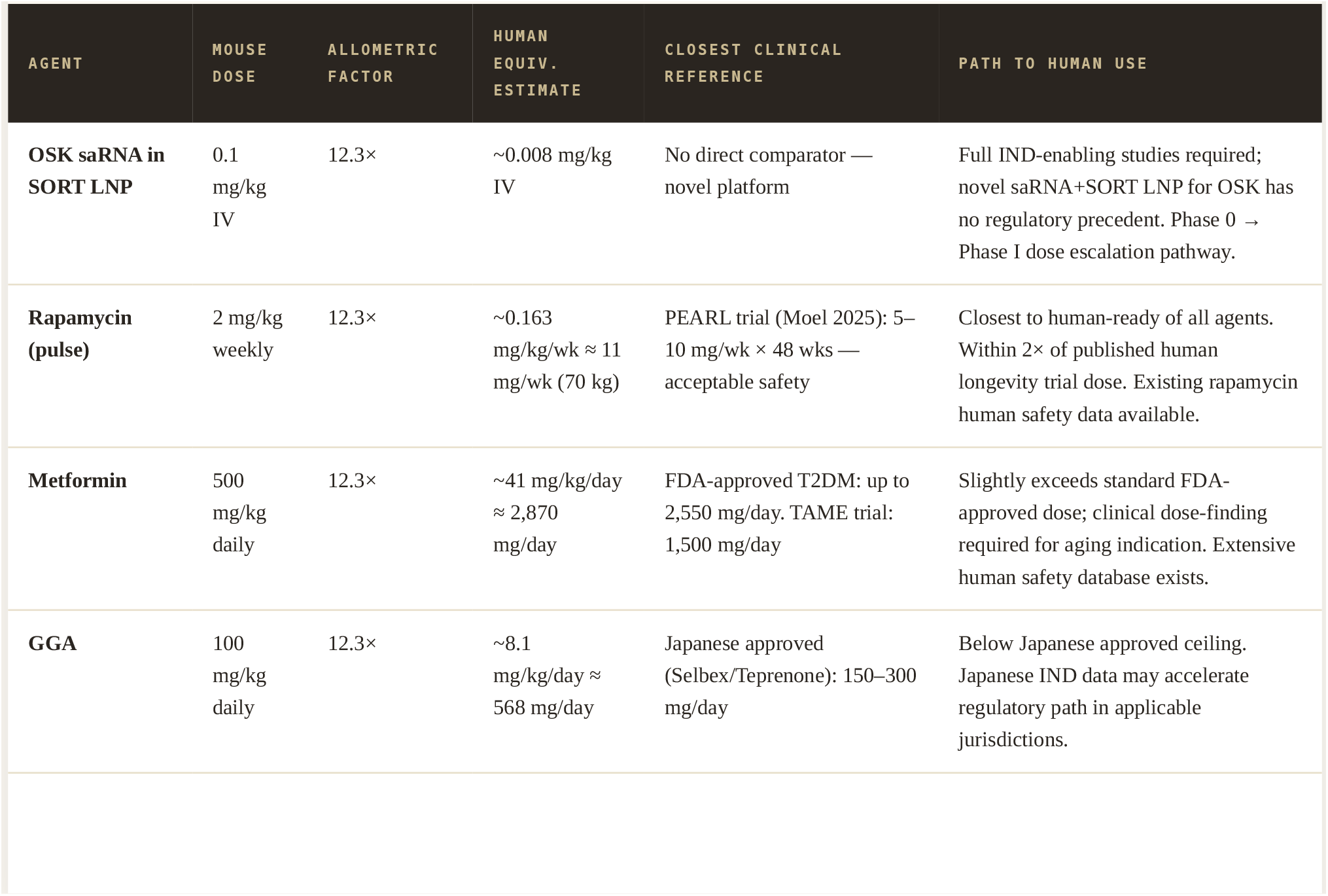
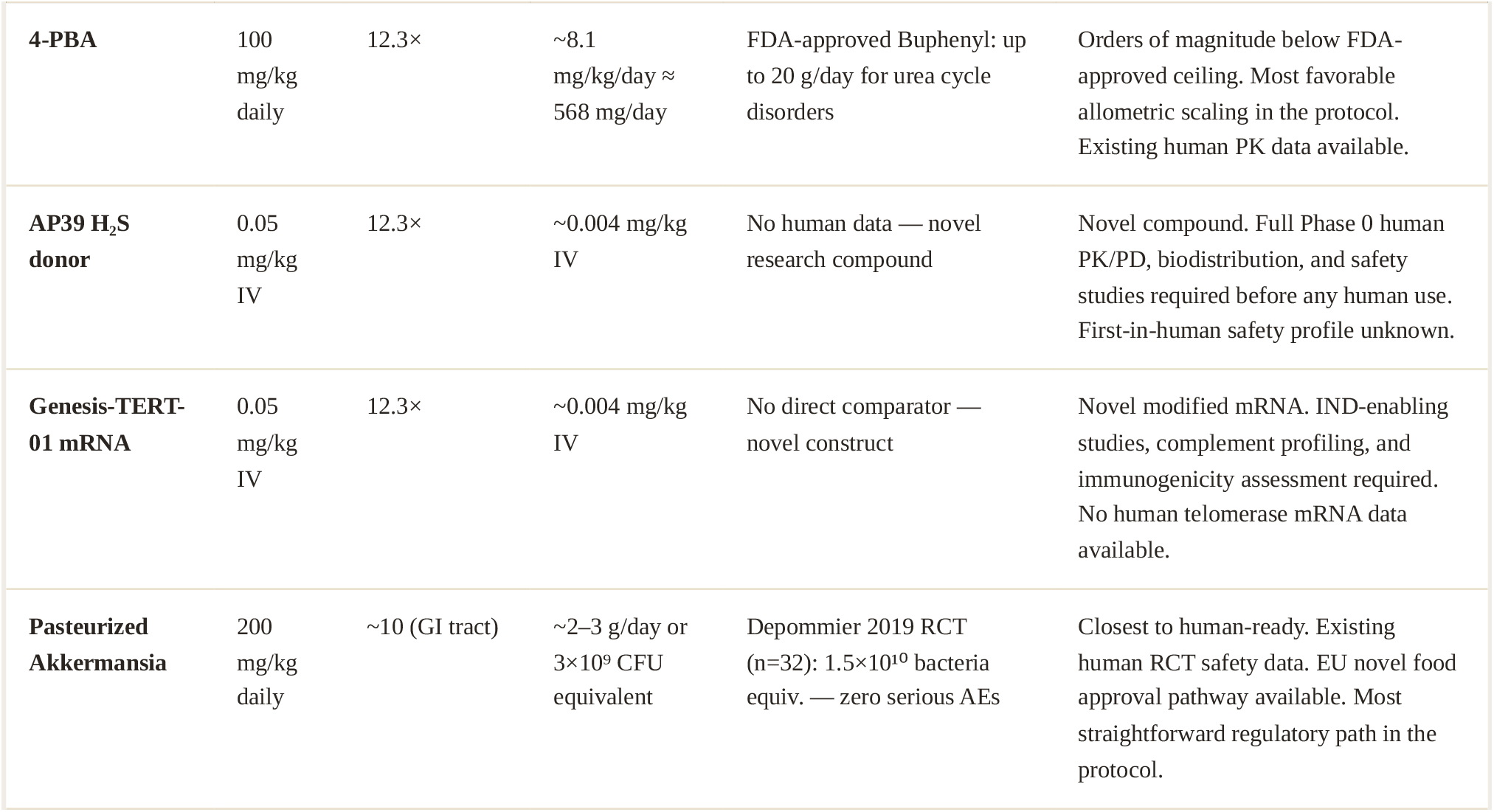

### FINAL DECLARATION Open Science Commitment & Final Declaration

The Cepeda Framework v1.1 is a broad, safety-oriented, open preclinical framework for testing whether coordinated partial rejuvenation strategies can be studied in a modular and falsifiable manner. **It is not a validated therapy. It is not a human protocol. It is a hypothesis-generating scientific blueprint ready for immediate preclinical testing**.

The framework accurately addresses all 12 official López-Otín 2023 aging hallmarks through an integrated non-integrating, open-source, CC0-licensed protocol. This claim is verified against the primary source. The framework additionally addresses a 13th Copenhagen 2022 candidate hallmark (RNA splicing dysregulation) through Companion C, pending independent HDF validation.

All citations in this document have been verified as real publications with accurate bibliographic data. All quantitative predictions are explicitly labeled as hypothesis-guiding model outputs requiring wet-lab falsification. All safety concerns identified through three cycles of independent review have been addressed with explicit protocol responses. The correction history is permanently documented in the scientific record.

**Open Science Commitment:** We commit to releasing all simulation code, model files, and output archives under CC0 at or before preprint submission. We commit to publishing all experimental results — positive, negative, and inconclusive — in real time on the public repository. Scientific credibility requires that failure is reported as diligently as success. We invite every qualified preclinical research laboratory worldwide to replicate, challenge, and improve this work. **The Cepeda Framework belongs to the scientific community**.

**CEPEDA FRAMEWORK CORE CONSTANTS ( GOSR-2026 ):**

OSK Ratio: OCT4:SOX2:KLF4 = 3:2:1 | Safeguards: 9 (parallel and sequential) | Kill-switch: LEAD-7294 dual miRNA gate |

Delivery: m1Ψ-saRNA + SORT LNP (5 mol% DOTAP) | Telomere: Genesis-TERT-01, 3-pulse (Days 7/9/11) | Companions: Pulse Rapamycin + Metformin + AP39 [HIGH-UNCERTAINTY; Tier 1A required] + GGA/4-PBA + Akkermansia/Synbiotic/Butyrate |

Multi-clock: Horvath + GrimAge2 + DunedinPACE + PhenoAge (07:00–09:00 standardized) |

Official hallmarks: 12/12 López-Otín 2023 + 1 Copenhagen 2022 candidate | Zero-NANOG Standard: <0.01% at all timepoints |

Simulation archive: 21,000 BioNetGen + Python stochastic cycles | Code: CC0 GitHub release at/before submission

### The Cepeda Framework v1.1 — Ready for Peer Review & Preclinical Validation

**Principal Investigator:** Cesar Cepeda — Executive Director, FTC Environmental Scientific Research Laboratories

FTC ENVIRONMENTAL SCIENTIFIC RESEARCH LABORATORIES · FORWARD THINKING COMMUNITIES · GOSR - 2026 · CC0 OPEN SCIENCE·

